# *Clostridioides difficile*-mucus interactions encompass shifts in gene expression, metabolism, and biofilm formation

**DOI:** 10.1101/2024.02.01.578425

**Authors:** Kathleen L. Furtado, Lucas Plott, Matthew Markovetz, Deborah Powers, Hao Wang, David B. Hill, Jason Papin, Nancy L. Allbritton, Rita Tamayo

## Abstract

In a healthy colon, the stratified mucus layer serves as a crucial innate immune barrier to protect the epithelium from microbes. Mucins are complex glycoproteins that serve as a nutrient source for resident microflora and can be exploited by pathogens. We aimed to understand how the intestinal pathogen, *Clostridioides diffiicile*, independently uses or manipulates mucus to its benefit, without contributions from members of the microbiota. Using a 2-D primary human intestinal epithelial cell model to generate physiologic mucus, we assessed *C. difficile-*mucus interactions through growth assays, RNA-Seq, biophysical characterization of mucus, and contextualized metabolic modeling. We found that host-derived mucus promotes *C. difficile* growth both *in vitro* and in an infection model. RNA-Seq revealed significant upregulation of genes related to central metabolism in response to mucus, including genes involved in sugar uptake, the Wood-Ljungdahl pathway, and the glycine cleavage system. In addition, we identified differential expression of genes related to sensing and transcriptional control. Analysis of mutants with deletions in highly upregulated genes reflected the complexity of *C. difficile*-mucus interactions, with potential interplay between sensing and growth. Mucus also stimulated biofilm formation *in vitro*, which may in turn alter viscoelastic properties of mucus. Context-specific metabolic modeling confirmed differential metabolism and predicted importance of enzymes related to serine and glycine catabolism with mucus. Subsequent growth experiments supported these findings, indicating mucus is an important source of serine. Our results better define responses of *C. difficile* to human gastrointestinal mucus and highlight a flexibility in metabolism that may influence pathogenesis.

**IMPORTANCE:** *Clostridioides difficile* results in upwards of 250,000 infections and 12,000 deaths annually in the United States. Community-acquired infections continue to rise and recurrent disease is common, emphasizing a vital need to understand *C. difficile* pathogenesis. *C. difficile* undoubtedly interacts with colonic mucus, but the extent to which the pathogen can independently respond to and take advantage of this niche has not been explored extensively. Moreover, the metabolic complexity of *C. difficile* remains poorly understood, but likely impacts its capacity to grow and persist in the host. Here, we demonstrate that *C. difficile* uses native colonic mucus for growth, indicating *C. difficile* possesses mechanisms to exploit the mucosal niche. Furthermore, mucus induces metabolic shifts and biofilm formation in *C. difficile*, which has potential ramifications for intestinal colonization. Overall, our work is crucial to better understand dynamics of *C. difficile*-mucus interactions in the context of the human gut.

## INTRODUCTION

As the leading cause of hospital-acquired diarrhea, *Clostridioides difficile* remains an urgent public health threat^1^. Although typically classified as a nosocomial pathogen, community-acquired cases of *C. difficile* infection (CDI) now comprise almost half the total number of cases^1^. As recurrent CDI affects nearly 50% of first-time patients^2^, a better mechanistic understanding of *C. difficile* pathogenesis is crucial to breaking this debilitating cycle. As an obligate anaerobe transmitted via spores, *C. difficile* germinates within the small intestine and establishes infection in the colon. *C. difficile* possesses mechanisms to directly adhere to the epithelium during colonization, including surface layer proteins^3,4^, flagella^5^, type IV pili^6^, and binary toxin in certain epidemic isolates^7–9^. Before accessing the epithelium, *C. difficile* must interact with colonic mucus, a key feature of innate host immunity.

The colonic mucus barrier is stratified, consisting of a diffuse luminal layer of secreted mucins inhabited by commensal microbes and a relatively sterile layer of membrane-bound mucins. The predominant secreted mucin in the colon is MUC2. Among membrane-bound mucins, MUC1 is particularly important in protection from bacterial invasion^10^. Past work suggests *C difficile* associates with the mucus barrier at multiple levels. Studies using animal colonization models indicate *C. difficile* inhabits the outer mucus layer^11^, while others showed co-localization of *C. difficile* and mucus in CDI patient stool samples, which are particularly rich in MUC1^12^. Subsequent work demonstrated direct adherence of *C. difficile* to purified MUC2^13^. This evidence indicates mucus serves as an anchoring point for *C. difficile* during colonization. Mucins are heavily glycosylated, with O-glycans decorating both MUC1 and MUC2 and N-glycans present on MUC1^10,14,15^. Glycans contribute to 80% of the mass of MUC2, and thus make up a significant proportion of mucin^16^. These glycans can be degraded by several bacterial species, providing a rich source of carbohydrates to the microbiota. Monosaccharides available from colonic mucins include fucose, mannose, galactose, N-acetylglucosamine (GlcNAc), N-acetylgalactosamine, and N-acetylneuraminic acid^14,15^. Following glycan cleavage, the peptide backbone also provides nutrients to the microbiota^14^. These backbones are rich in serine, threonine, and proline^14,17^. In humans, nearly 45% of the amino acid composition of MUC1 and over 55% of MUC2 consists of serine, threonine, and proline based on canonical sequences in UniProt (P15941 and Q02817, respectively); others have predicted greater proportions of these amino acids^18^. Overall, interactions between commensal microbiota and mucus are often symbiotic, resulting in a thicker, more protective mucus layer for the host and increased nutrient availability for the microbiota^14,19^. To maintain healthy conditions, mucin degradation by bacteria and regeneration by the host must be carefully balanced.

Pathogens can alter and exploit mucus during infection^10^. Previous work showed that oligosaccharide composition within mucus is altered during CDI, and that *C. difficile* reduces expression of human *MUC2* while preferentially interacting with MUC1^12^. *C. difficile* also benefits from the cleavage of mucins by specific members of the microbiota^20^, however, the extent to which mucus is altered or metabolized specifically by *C. difficile* remains unclear. The carbohydrate active enzymes (CAZy) database indicates *C. difficile* R20291 possesses enzymes from 31 families^21^, at least one of which, glycosyl hydrolase family 38, contains enzymes likely involved in mucin degradation^20^. To our knowledge, no study has assessed the mucolytic capacity of any enzymes listed in the CAZy database for *C. difficile*. Nonetheless, evidence to date indicates that alterations to mucus promote *C. difficile* colonization.

There are substantial challenges in obtaining mucus that accurately recapitulates native human mucus in its composition and viscoelasticity^22^, properties likely vital to *C. difficile*-mucus interactions. Human mucus varies from that of animal models, and the processing of commercial mucins removes important components from native mucus and disrupts its structure^23^. Furthermore, immortalized colonic cell lines often do not secrete the same proportions of mucin types as those in a healthy colon, if mucins are secreted at all^23–25^. Recently, a human primary intestinal epithelial cell (IEC) co-culture system was validated for use with *C. difficile*^26^. These IECs can secrete a thick mucus barrier^27^, which can be harvested or directly inoculated to assess *C. difficile*-mucus interactions. Importantly, the biophysical properties and composition of IEC-derived mucus are similar to mucus derived from *ex vivo* human tissues^28^.

The goal of this study was to assess specific interactions between *C. difficile* and human colonic mucus to better understand the extent to which *C. difficile* can independently use or manipulate mucus to its benefit. Using the physiologically relevant, primary IEC-derived mucus described above, we measured the contribution of mucus to *C. difficile* growth. We then used transcriptomics to explore the response of *C. difficile* to mucus and used these data in metabolic modeling to predict how mucus shapes *C. difficile* metabolism. We additionally assessed the capacity of *C. difficile* to alter biophysical and biochemical properties of mucus. Our work provides a multi-faceted understanding of *C. difficile*-mucus interactions, which may influence colonization or disease progression.

## RESULTS

### Mucus derived from primary human IECs promotes *C. difficile* growth

To test whether colonic mucus derived from primary human IECs supports or promotes *C. difficile* growth, we examined growth *in vitro* in *C. difficile* minimal medium (CDMM)^29^ containing 1% glucose with and without 50 µg/mL mucus. We also tested conditions without glucose, wherein mucus was the only source of sugars. All media tested supported *C. difficile* growth due to the presence of casamino acids. In the presence of mucus, *C. difficile* exited lag phase earlier and reached higher maximum OD_600_ than without mucus, both with and without glucose (Fig 1A).

**Figure 1.**
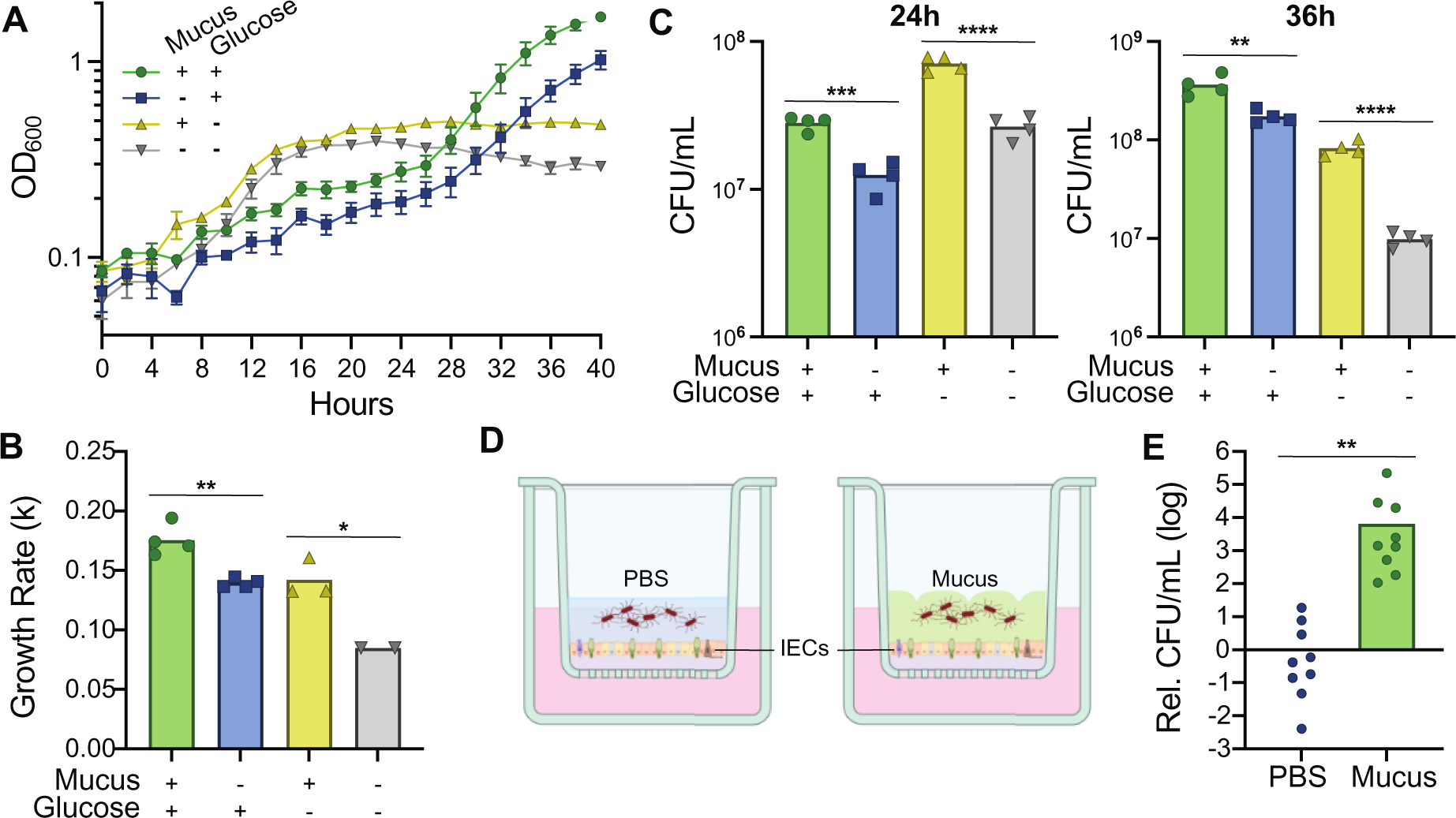
Mucus derived from primary human IECs enhances *C. difficile* growth. **(A)** *C. difficile* R20291 growth curves in CDMM containing purified mucus and glucose (+/+), glucose only (-/+), purified mucus only (+/-), or no mucus or glucose (-/-). Data are from one representative experiment, n=4. **(B)** Growth rates during exponential phase in each medium. Exponential phase was defined by having at least three time points in a linear range; samples for which at least three time points in a linear range could not be identified were excluded. **(C)** Viable cell counts expressed as CFU/mL after 24 hours (left) and 36 hours (right) growth in each medium. **(D)** Schematic of the primary human IEC co-culture system. IECs secrete produce a thick mucus barrier that can be inoculated with bacteria (right). As a control, the mucus layer was removed mechanically and replaced with PBS (left). **(E)** *C. difficile* viable cell counts after 2h co-culture with IECs. CFU/mL values were normalized to the CFU/mL present in the respective inoculum, then expressed relative to the mean normalized CFU/mL in the PBS condition. Data are from three independent experiments, n=3 per experiment. *p < 0.05, **p < 0.01, ***p < 0.001, ****p < 0.0001, unpaired, two-tailed t-test.

In CDMM with glucose, the addition of mucus resulted in faster growth rates (*k*) during exponential phase (*k* = 0.176±0.013 OD_600_/hour) compared to the no-mucus condition (*k* = 0.140±0.004 OD_600_/hour) (Fig 1B). At 24h, cultures with glucose and mucus contained 2.25-fold more CFU/mL on average than cultures without mucus (Fig. 1C). By the end of exponential phase at 36h, we again recovered more viable cells from conditions with mucus, with 2.10-fold more CFU/mL than without mucus. Without glucose, *C. difficile* exhibited a higher growth rate during exponential phase with mucus (*k* = 0.142 ± 0.085 OD_600_/hour) compared to the no-mucus condition (*k* = 0.085±0.000 OD_600_/hour) (Fig 1B). In addition, the presence of mucus increased the number of viable cells; we recovered on average 2.68-fold more CFU/mL at 24h and 8.43-fold more CFU/mL at 36h from cultures grown with mucus versus without (Fig. 1C). Altogether, differences in optical density, growth rates, and CFU indicate mucus enhances *C. difficile* growth, both when mucus supplements glucose and when mucus is the sole carbohydrate source.

Because *C. difficile* exhibited increased growth in media with IEC-derived mucus, we examined growth in co-culture with IECs with and without an intact mucus layer. The IECs were stimulated to produce a robust mucus layer during differentiation as previously described (Fig. 1D, right)^27^. For controls without mucus, we removed the mucus layer and replaced it with PBS (Fig. 1D, left). *C. difficile* was then inoculated at an MOI ∼0.01. After 2h in co-culture, we observed 14.0-fold greater expansion in CFU in co-cultures where the mucus layer was intact compared to those without mucus (Fig. 1E). Overall, our results indicate that IEC-derived mucus promotes *C. difficile* growth, both in broth and in the context of an infection model.

### Transcriptional profiling suggests mucus alters expression of genes with roles in metabolism and nutrient acquisition

To determine how *C. difficile* responds to the IEC-derived mucus, we used RNA-seq to assess transcription in exponential-phase cultures grown with and without 50 µg/mL mucus in CDMM. Because we were most interested in responses to mucus as opposed to stress or starvation, 1% glucose was retained in the media. We identified 282 upregulated and 285 downregulated genes in the presence of mucus (567 total; fold change > 2, p-adj < 0.05) (File S2, Fig. 2A). Among the five genes most upregulated in mucus, three are potential transporters (CDR0455, CDR1626, and CDR2495), one is annotated as a TetR transcriptional regulator (CDR0508), and one is annotated as a putative xanthine/uracil permease (CDR2014). The five most downregulated genes are largely annotated as putative or hypothetical proteins, with one probable protease (CDR3145). Principal components analysis indicated that mucus was the main variable contributing to variance in the dataset (Figure 2B).

**Figure 2.**
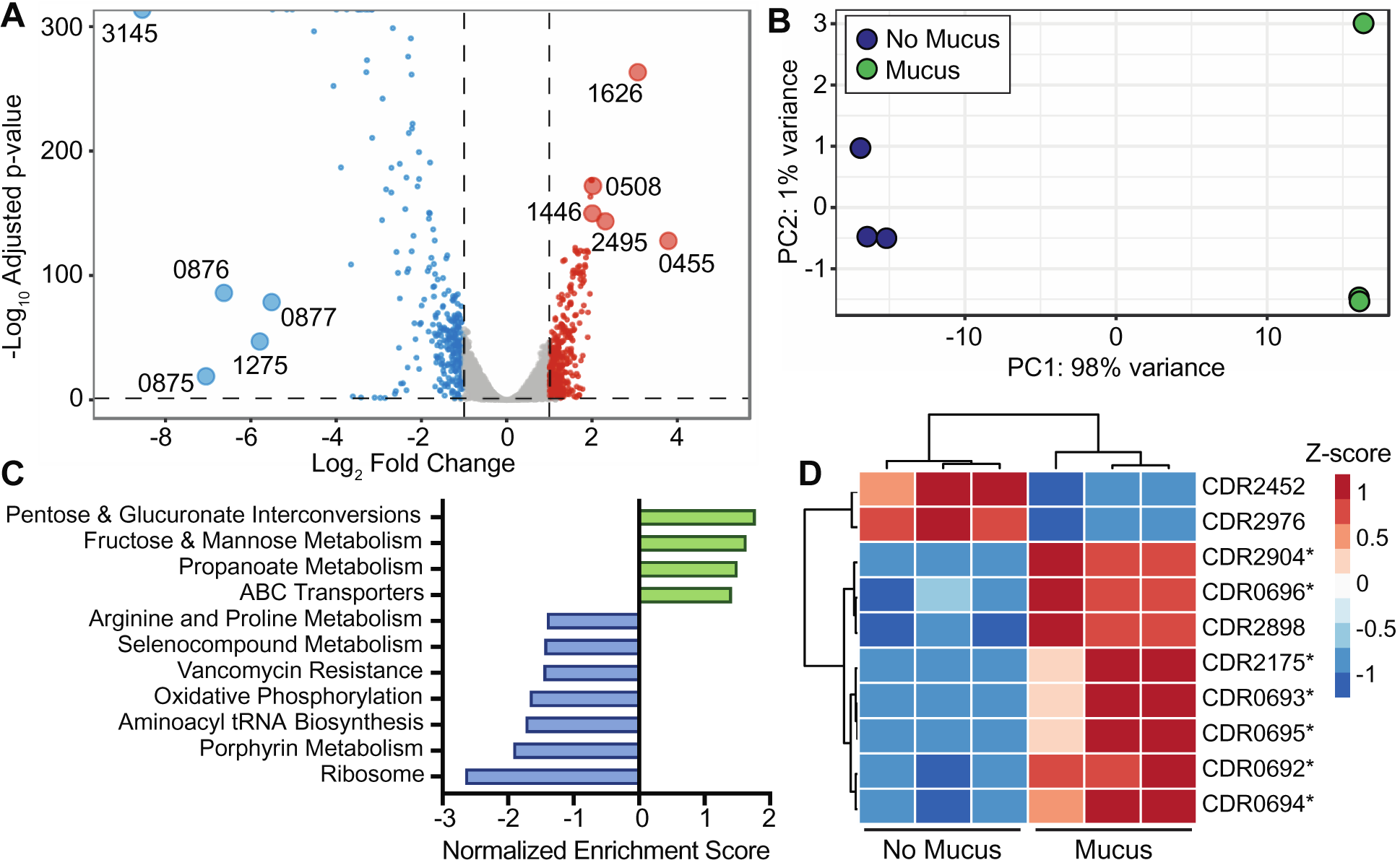
Mucus leads to a distinct and robust transcriptional response affecting metabolism. **(A)** Volcano plot highlighting all genes differentially expressed in CDMM with mucus vs. no mucus with Benjamini-Hochberg adjusted p-value < 0.05 and fold changes > 2 (log_2_ fold changes > 1 or < −1), based on Wald test in DESeq2. Genes significantly upregulated (red) and downregulated (blue) with mucus are shown. The 5 most upregulated and downregulated genes are labelled by the last 4 digits of their locus tag (CDR20291_XXXX, hereafter abbreviated as CDRXXXX). **(B)** PCA plot after variance stabilizing transformation of read counts in DESeq2, n=3 per condition. **(C)** Normalized enrichment scores for KEGG gene sets identified using GSEAPreranked considering relative expression all genes in *C. difficile* R20291. The selected gene sets were enriched in the presence (green) or absence (blue) of mucus, nominal p < 0.05 and/or FDR q-value < 0.25. **(D)** Heatmap of genes in the fructose and mannose metabolism gene set identified using GSEAPreranked based on genes with adjusted p < 0.05 and fold change > 2 from RNA-Seq. Starred genes contributed to core enrichment of the gene set.

We next used gene set enrichment analysis (GSEA) to identify KEGG pathways enriched in each condition. Using GSEAPreranked with all genes in *C. difficile*, we identified four gene sets enriched with mucus and seven enriched without mucus (Fig. 2C, File S3). We further refined these results using GSEAPreranked with the 567 most differentially expressed genes. This analysis identified no gene sets significantly enriched in conditions without mucus; with mucus, only the fructose and mannose metabolism pathway remained enriched (nominal p = 0.049, FDR q = 0.197). Figure 2D shows relative expression of highly differentially expressed genes in this gene set. Among the seven genes contributing to core enrichment, five are annotated as phosphotransferase system (PTS) components, which are important for sugar uptake (Table S1). Specifically, CDR0692-0696 are involved in the transport of sorbitol, a host- and diet-derived metabolite^30^, while CDR2904 likely encodes part of a mannose PTS. These analyses suggest mucus alters sugar uptake and metabolism in *C. difficile*, which prompted us to search for additional genes that could play a role in utilizing nutrients from mucus.

Increased proportions of amino acids in the gut can lead to increased susceptibility for CDI^31^, with Stickland metabolism playing a key role in the conversion of amino acids for growth^32^. Proline and glycine are particularly important for reductive Stickland metabolism, in which proline and glycine reductases (PR and GR, respectively) are used to regenerate NAD+^33^. Our analysis showed that all but three of the 19 genes within PR and GR clusters were significantly downregulated with mucus (Table S1), suggesting that Stickland reduction of proline and glycine is suppressed by *C. difficile* during mid-exponential phase when mucus is present.

Recent work has revealed the importance of the Wood-Ljungdahl pathway (WLP), which provides metabolic flexibility to *C. difficile* and potential advantages during infection^34,35^. Expression of WLP genes tends to be inversely correlated with PR and GR^36^, and increased expression of WLP genes could indicate an abundance of Stickland electron donors, such as alanine, valine, serine, isoleucine, threonine, or glutamic acid^36,37^. Under such conditions, alternative mechanisms of reduction are needed^36^. Given apparent inhibition of PR and GR by mucus, we investigated WLP gene expression. Of 38 genes that have been previously identified as part of the WLP or a linked glycine cleavage system (GCS, explained below) in *C. difficile*, 19 were upregulated and 15 were downregulated with mucus (p-adj < 0.05, File S2). We observed increased expression of genes corresponding to branches of the WLP that fix CO_2_ and convert it to acetyl-CoA (Table S1). Upregulated genes encoding enzymes in these branches included: *fdh* and *hyd* genes for reversible conversion between formate and CO_2_; *metV* and *metF* encoding components of N^5^,N^10^-methylene-tetrahydrofolate reductase, *cooS* for fixation of CO_2_ via CO dehydrogenase; and homologs of *acsE, acsC, acsD*, and *acsB*, which encode components of acetyl-CoA synthase. WLP genes downregulated in mucus corresponded to interconversions between acetyl-CoA and acetate, butyrate, or ethanol. These included homologs of *pta* and *ptb*, encoding enzymes that convert between acetyl-CoA and acetylphosphate, as well as *ackA* and *buk* for conversion between acetylphosphate and acetate. We also observed downregulation of *thlA, hbd, crt2,* and *bcd-etfAB* homologs for conversion of acetyl-CoA to butyrate, and of *adhE*, for conversion of acetyl-CoA to ethanol via acetaldehyde.

Linked to the WLP is a reversible glycine cleavage system (GCS) that provides additional options for carbon assimilation in *C. difficile*^36,38^. We observed increased expression of *gcvH* and *gcvL*, and decreased expression of *gcvP* and *gcvT*, which encode components of the glycine cleavage reaction complex^39,40^ (Table S1). Expression of *glyA,* which encodes a serine/glycine hydroxymethyltransferase to interconvert glycine and serine, was also increased. In addition, *sdaB,* which encodes a serine dehydratase that produces pyruvate from serine, was upregulated. Overall, the presence of mucus decreased expression of PR and GR gene clusters, but increased expression of genes related to CO_2_ fixation via the WLP and glycine and serine catabolism via the GCS.

### Mechanisms for transcriptional control are differentially expressed with mucus

An abundance of genes for transcriptional regulation and responding to environmental stimuli were differentially expressed (File S2), suggesting *C. difficile* senses mucus. Of highly differentially expressed genes with fold change > 2 (Table S2), many are annotated to encode transcriptional regulators, sigma factors, or antiterminators. Genes encoding transcriptional regulators from the GntR family were most prevalent, and genes annotated as members of the TetR, MarR, AraC, and MerR regulator families were also differentially expressed. These families can regulate many cellular processes, including overall metabolism of carbon and nitrogen (GntR, AraC), stress responses (TetR, AraC), and resistance to antibiotics, metals, or other toxins (MarR, MerR)^41^.

Two-component systems play a critical role in sensing and responding to environmental stimuli. Among TCS genes (Table S2), CDR1568-1569 were upregulated, while CDR2206-2205, *hexRK*, CDR2021-2020, and CDR2188-2187 were downregulated. Based on work in *C. difficile* and with orthologous genes in *B. subtilis*, CDR1568-1569 could be involved in maintaining cell surface homeostasis^42^, while *hexRK*, and potentially CDR2021-2020, are involved in antibiotic sensing and resistance^43,44^. Altogether, our data suggest mucus is an important stimulus for *C. difficile,* perhaps indicating increased nutrient availability or proximity to the epithelium.

### Mutants lacking upregulated genes exhibited altered growth phenotypes with mucus

Among the genes most highly upregulated in *C. difficile* grown with mucus, we identified several predicted to be involved in transport. CDR0455 and CDR2495 were among the five most upregulated genes in mucus (Fig. 2A). CDR0455 forms a predicted operon with CDR0454-0453, which were also significantly upregulated (Fig. 3A, File S2). Based on analyses of protein homology in Phyre2^45^, the operon likely encodes three membrane proteins consisting of an endopeptidase (CDR0453), Na^+^/H^+^ antiporter (CDR0454), and transport protein (CDR0455).

**Figure 3.**
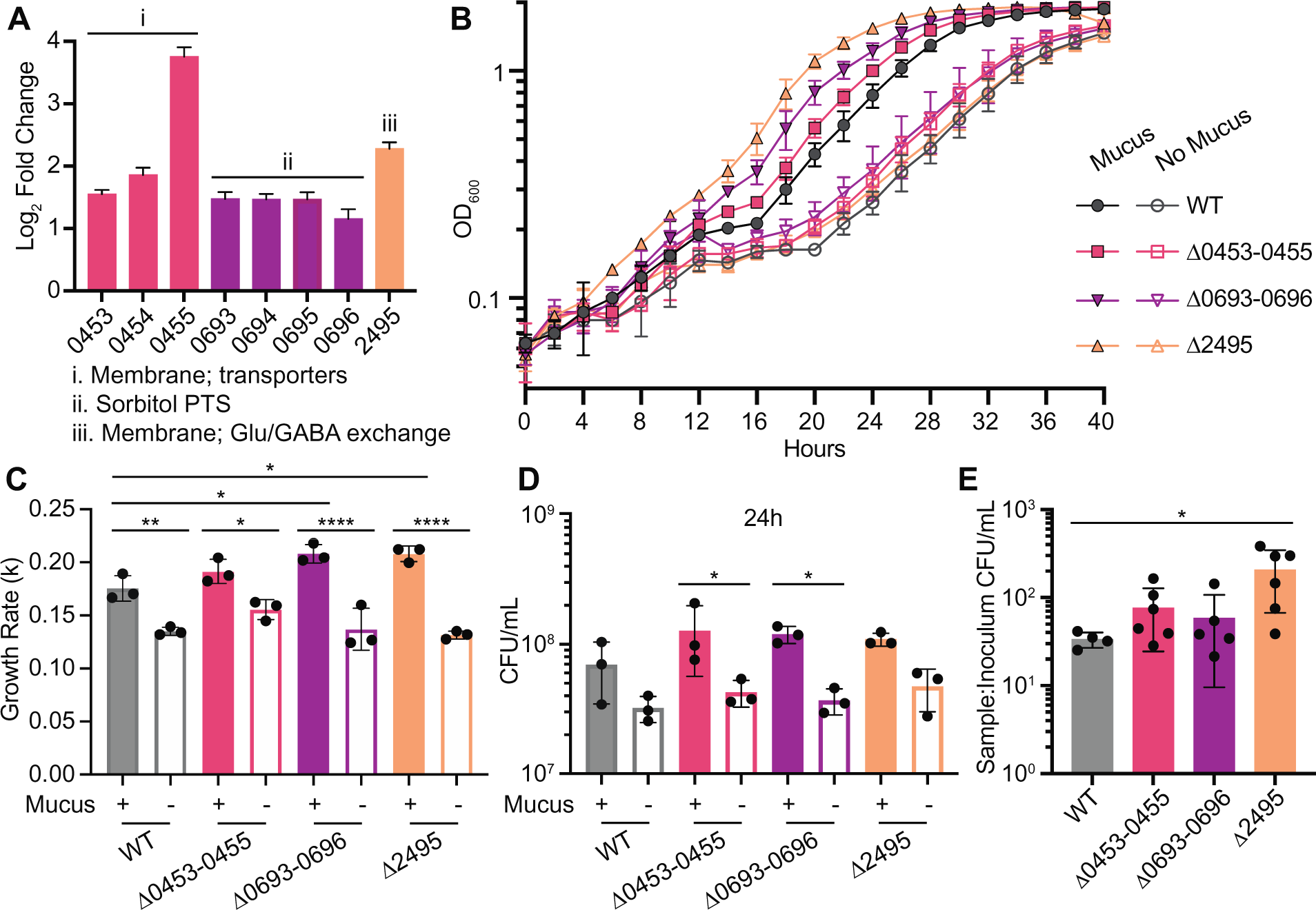
Deletion of genes and operons upregulated with mucus enhances growth. **(A)** Log_2_ fold change expression values and standard error for genes and operons selected for mutagenesis. CDR0453-0455 and CDR2495 were among the most highly upregulated; CDR0693-0696 contributed to core enrichment of the fructose and mannose metabolism gene set in mucus (Fig 2). Predicted functions based on Phyre2 analysis or prior studies are indicated. **(B)** Growth curves for mutants and wildtype in CDMM with mucus (filled symbols) and without mucus (open symbols). Data are from one representative experiment, n=3. **(C)** Growth rates during exponential phase. **(D)** Viable cell counts expressed as CFU/mL. **(E)** Viable cell counts after 6h co-culture with IECs with an intact mucus barrier. Viable cell counts are expressed as CFU/mL in each sample after 6h, normalized to CFU/mL present in respective inoculum for each strain. Data are combined from two independent experiments, n=2 or 3 per experiment. Outliers were determined and removed using Grubbs’ method. *p < 0.05, **p < 0.01, ***p < 0.001, ****p < 0.0001, one-way ANOVA with Tukey’s or Sidak’s tests

CDR2495 encodes a membrane protein with homology to *E. coli* GadC, indicating potential function in glutamate/γ-aminobutyrate exchange. GSEA identified upregulated PTS components within the fructose and mannose metabolism gene set (Figure 2C, 2D, Table S1). Several of these genes belong to a predicted operon encoding a sorbitol PTS encompassing *gutM, gutA, srlE* (CDR0693)*, srlE’, srlB,* and *gutD*, all of which were significantly upregulated in mucus (Fig. 3A, Table S1). Using qRT-PCR, we independently confirmed increased expression of CDR0455, CDR0693, and CDR2495 under conditions used in the RNA-Seq experiment (Fig. S1). Given consistent upregulation of these genes and their potential role as transporters, we predicted that deletions in these genes would result in reduced growth in mucus. Thus, we generated in-frame gene and operon deletions in *C. difficile* R20291: Δ0453-0455, Δ0693-0696, and Δ2495.

We evaluated growth of the mutants in CDMM containing 1% glucose with or without 50 µg/mL mucus (Fig. 3B). Like wildtype, all mutants had higher growth rates with mucus than without (Fig. 3C). While wildtype and mutants showed similar growth in the absence of mucus, unexpectedly, the Δ0693-0696 and Δ2495 mutants had significantly higher growth rates relative to wildtype in mucus. Furthermore, after 24h Δ0453-0455 and Δ0693-0696 had 2.99-fold and 3.24-fold more CFU/mL with mucus, relative to respective growth without mucus (Fig. 3D). We observed similar, albeit statistically insignificant, trends with wildtype and Δ2495 in mucus. In media lacking glucose, we again observed that all strains had significantly more growth overall with mucus than without mucus based on OD_600_ measurements and CFU at 24h (Fig. S2A-C). However, in the absence of glucose, mucus did not significantly alter growth rates for any strain (Fig. S2D), and the increased growth rates and shorter lag phases of mutants in conditions with glucose were not observed.

To assess growth phenotypes in a native mucus layer, we evaluated the growth of each mutant in co-culture with IECs and an intact mucus layer. After 6h co-culture, we observed at least one log of growth for all strains (Fig. 3E). Consistent with results from broth culture experiments, the mutants tended toward more growth relative to wildtype, with significantly greater growth of Δ2495 (6.17-fold more CFU/mL for Δ2495 vs. wildtype; Fig. 3E). Altogether, mutations in the selected genes and operons did not lead to growth defects in mucus as we predicted, but instead resulted in greater growth than wildtype under these conditions.

### *C. difficile* biofilm formation may increase viscosity of *ex vivo* mucus

To evaluate the extent to which *C. difficile* can manipulate mucus, we performed biophysical and biochemical analyses on native *ex vivo* mucus incubated for 24 hours with and without *C. difficile*. Importantly, *C. difficile* remained viable and expanded within *ex vivo* mucus (Fig. S3A). We applied particle tracking microrheology (PTMR) to measure the viscosity *C. difficile*- and mock-inoculated mucus, using a Gaussian mixture model to distinguish watery (less viscous) vs. mucoid (more viscous) fractions of mucus^28,46–48^. After 24 hours, mucus containing *C. difficile* had more mucoid signal relative to the mock-inoculated control, as indicated by increased detection of microbeads within the mucoid vs. watery fraction (Fig. 4A, 4C). This increased mucoid fraction corresponded to greater complex viscosity in samples containing *C. difficile* relative to the mock-inoculated control (Fig. 4B). Multiangle laser light scattering (MALLS), which measures mucin molecular weight, radius of gyration, and concentration, indicated no changes in these metrics between inoculated and mock-inoculated samples after 24h (Fig S3B-D).

**Figure 4.**
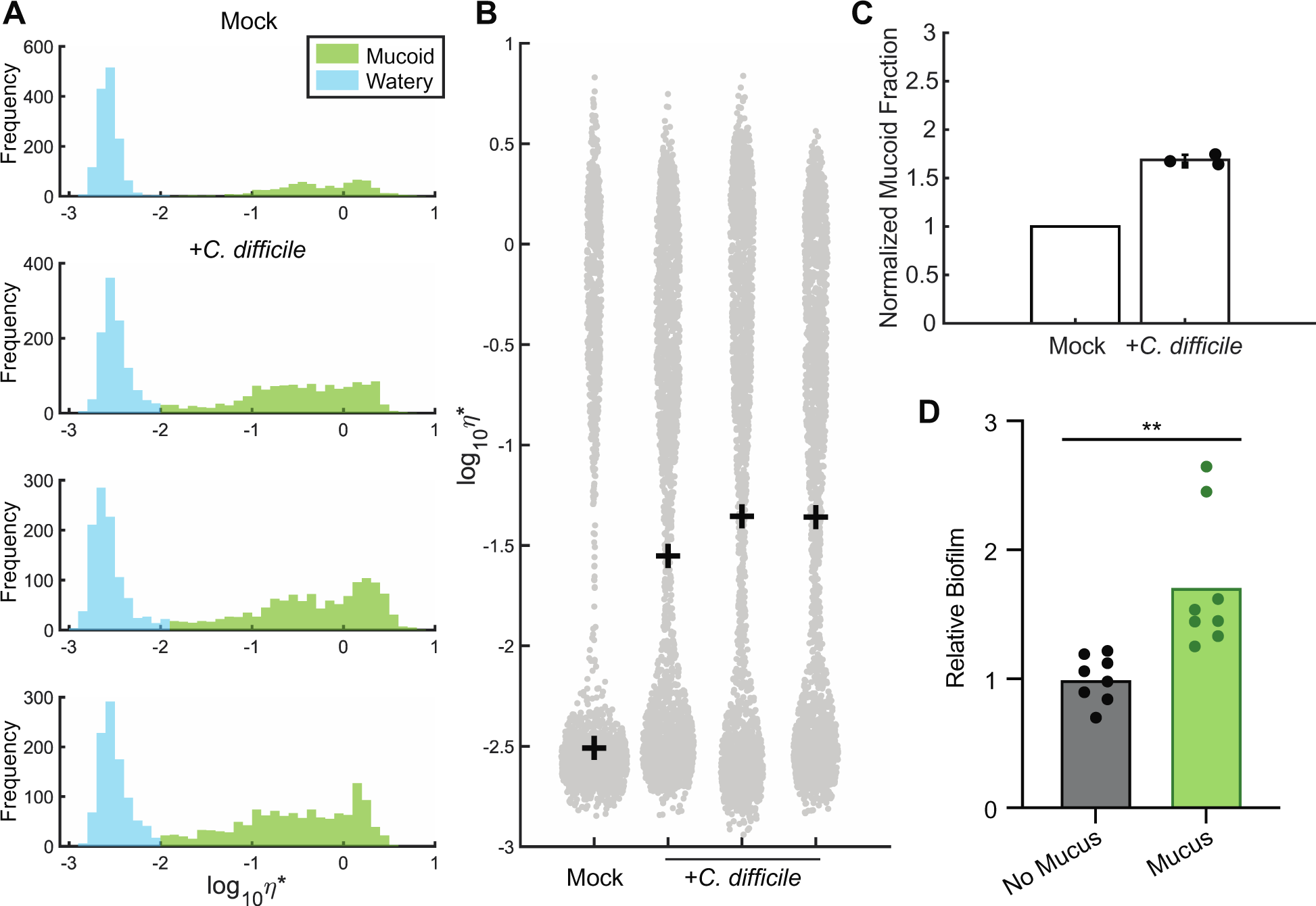
*C. difficile* increases viscosity of *ex vivo* mucus. **(A)** PTMR analysis of *ex vivo* IEC mucus inoculated with *C. difficile* or mock-inoculated with 1X CDMM salts. A Gaussian mixture model was employed to distinguish mucoid rheological behavior from that of watery mucus after 24 hours. An increasing frequency of detection of microbeads within the mucoid fraction is indicated by larger green peaks. **(B)** Complex viscosity (η*) of mucus-grown samples after 24h. Individual points represent measured complex viscosity from each microbead, and crosses represent mean η* per sample. **(C)** Quantified mucoid fractions in mucus inoculated with *C. difficile* relative to the mock-inoculated control based on PTMR data in (A) For each sample, three slides were prepared and 10 videos recorded per slide. **(D)** Biofilm formation by *C. difficile* in CDMM with or without mucus after 24h, expressed as absorbance at 570nm for each sample relative to mean absorbance of the no mucus condition. Data from 2 independent experiments, n=4 per experiment, were combined. **p < 0.01, unpaired two-tailed t-test.

While results indicate that *C. difficile* does not break down mucins sufficiently to detect decreased molecular weights, the observed increase in viscosity could reflect additional ways *C. difficile* responds to mucus. Biofilm formation within mucus can lead to increased viscosity, as observed with *Pseudomonas aeruginosa*^48^. We therefore examined the impact of mucus on *C. difficile* biofilm formation *in vitro*. We detected significantly greater biofilm biomass for *C. difficile* grown with mucus versus without mucus (Fig. 4D), indicating that mucus promotes biofilm formation, which may contribute to the increased viscosity we observed.

### Metabolic modeling predicts increased uptake of specific amino acids from mucus

To assess the metabolic potential of *C. difficile* during growth in mucus, we used RIPTiDe to contextualize an established genome-scale metabolic network reconstruction (GENRE) for *C. difficile* R20291 with our transcriptomic data^49–51^. To recapitulate conditions from the RNA-Seq experiment, we allowed the model to use mucin-derived monosaccharides for conditions with mucus and excluded them from the no-mucus condition. Ordination analyses revealed that predicted core metabolic activity was distinct between conditions with and without mucus (Fig. 5A). These differences in metabolic activity corresponded to a predicted 52.6% increase in biomass flux, a proxy for growth rate, in conditions with mucus versus without (Fig. 5B). Overall, comparing biomass fluxes to experimental data (Fig. 1, 3) suggests the model accurately predicted growth trends given respective transcriptomic contexts.

**Figure 5.**
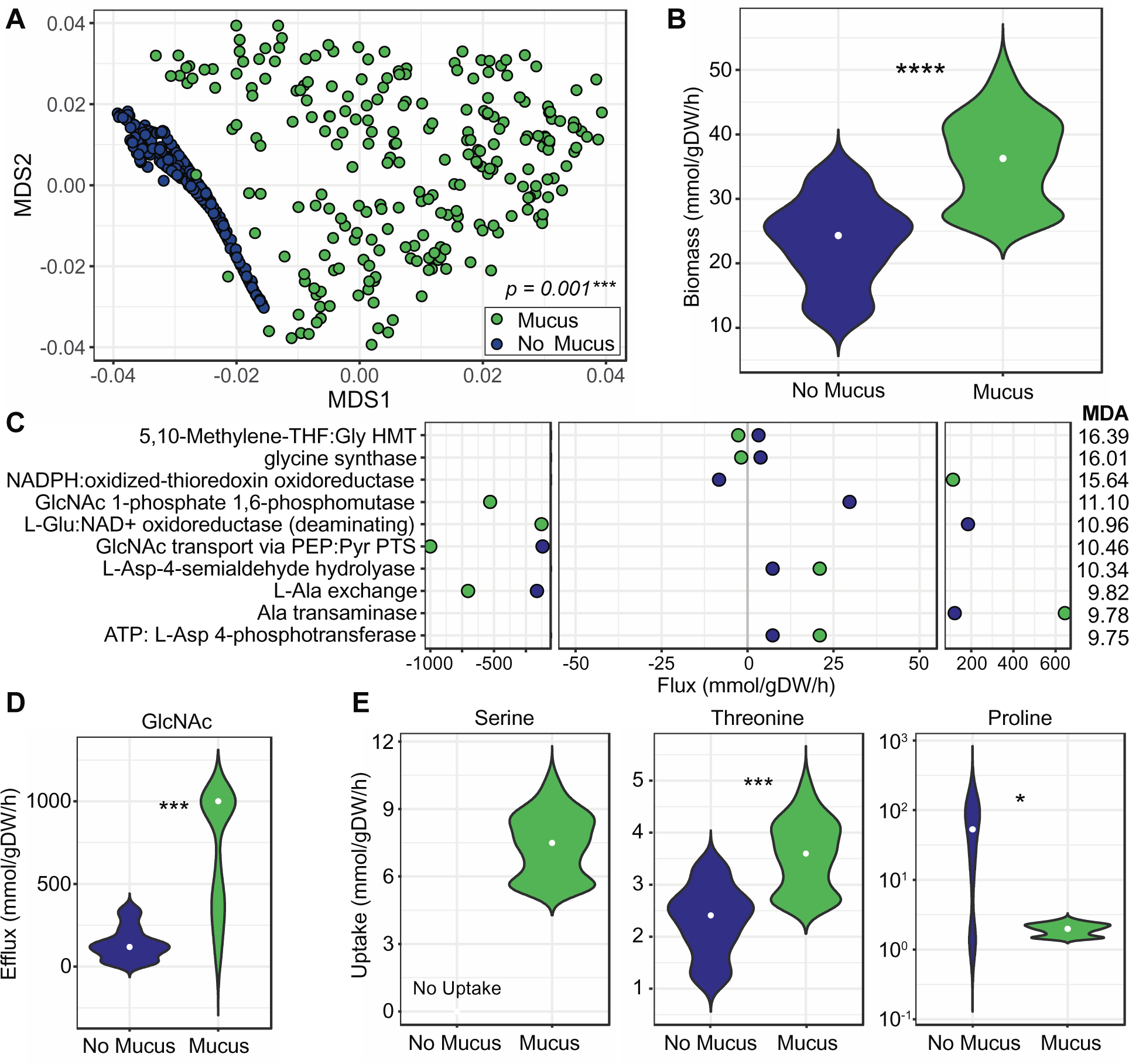
Modeled *C. difficile* growth predicts distinct metabolic activity in the presence of mucus. (A) Predicted core metabolic activity for *C. difficile* from conditions with vs. without mucus, using a *C. difficile* GENRE contextualized with transcriptomic data. Data are represented as NMDS ordination of the Bray-Curtis dissimilarity of flux distributions from shared reactions in each contextualized model. Differences between conditions were determined by PERMANOVA. (B) Predicted biomass flux in CDMM with vs. without mucus. (C) Median fluxes for reactions determined by Random Forests analysis to be important in differentiating conditions with (green) vs. without mucus (blue). Reactions are ranked by MDA with larger values indicating greater importance. (D) Predicted GlcNAc efflux in conditions with vs. without mucus. (E) Predicted uptake of the indicated amino acids in conditions vs. without mucus. Data were generated from n=250 samplings from each contextualized model. Median flux values in B, D, and E are indicated (white dot). Differences in B, D, and E were determined by measuring p-values from a Wilcoxon rank sum test based on n=12 random subsamples from each condition (5% of total samples). This random subsampling and testing process was repeated 1000 times and the median p-value was used: *p < 0.05, **p < 0.01, ***p < 0.001, ****p < 0.0001.

To rank relative importance of reactions in contributing to predicted differences in biomass, we used a Random Forests classifier to determine mean decrease accuracies (MDA) (Fig. 5C). The positive or negative median flux values shown indicate directionality of each reaction. The first two reactions, corresponding to glycine hydroxymethyltransferase (GHMT) and glycine synthase, are involved in the GCS. Negative flux values for GHMT indicated conversion from serine to glycine in the mucus condition, while flux in the opposite direction was predicted without mucus. Negative flux for glycine synthase also indicated biosynthesis of glycine from 5,10-methylenetetrahydrofolate, which simultaneously generates NAD+, in the presence of mucus. NADPH:oxidized-thioredoxin oxidoreductase and L-Glu:NAD+ oxidoreductase are also involved in redox chemistry. In the presence of mucus, positive flux indicated oxidation of NADPH and reduction of thioredoxin was predicted for the former, and negative flux indicated oxidation of NADH and reduction of 2-oxoglutarate to L-glutamate was predicted for the latter. These results suggest glycine biosynthesis and regeneration of electron carriers is important in the presence of mucus.

Reactions converting or transporting GlcNAc were also among the most important. GlcNAc is a mucin-derived monosaccharide that can stimulate biofilm formation in *C. difficile*^52^, and is a component of biofilms^50,53^. For GlcNAc-1-phosphate 1,6-phosphomutase, negative flux values with mucus indicated conversion from GlcNAc-1-phosphate to GlcNAc-6-phosphate (Fig. 5C). For GlcNAc transport, negative flux values, which were 8.42-fold greater with mucus than without, indicated conversion of GlcNAc-6-phosphate to GlcNAc. Direct analysis of GlcNAc exchange suggested that GlcNAc efflux was due to flux through the above reactions, with 5.16-fold greater efflux with mucus than without (Fig. 5D).

Because glycine and serine are involved in reactions with the greatest MDA, we examined predicted uptake for these amino acids. Glycine exchange was predicted to be negligible in both conditions. Serine uptake, however, was predicted only in conditions with mucus (Fig. 5E). This result makes sense considering flux through GHMT predicted biosynthesis of glycine from serine in conditions with mucus, whereas without mucus, opposite flux toward serine biosynthesis was predicted. As mucins are proline-, threonine-, and serine-rich, we also examined predicted uptake of proline and threonine. We observed 1.53-fold greater threonine uptake with mucus versus without. However, we observed the opposite trend for proline uptake, with 29.4-fold more uptake predicted for the no-mucus condition versus with mucus. These results coincide with RNA-Seq data suggesting proline metabolism via PR is less active with mucus. Nonetheless, predicted increases in threonine and serine uptake in mucus are intriguing as glycans are attached to mucins at serine and threonine residues.

### Mucus restores growth in defined media lacking specific amino acids

Metabolic modeling predicted glycine and serine interconversion to be particularly important for differentiating growth with versus without mucus, and also predicted greater uptake of serine and threonine from conditions with mucus. Hence, we experimentally investigated the importance of these amino acids using DCAMM, a modified version of CDMM with defined amino acid composition. We assessed growth with and without mucus in media containing all amino acids (complete), lacking threonine (-T), or lacking serine (-S). Due to interconversion of glycine and serine, we also assessed media lacking glycine (-G) or lacking glycine and serine (-G-S). Importantly, complete DCAMM supported *C. difficile* growth comparably to CDMM, with similar patterns of growth, including higher growth rates and more CFU/mL at exponential phase, with versus without mucus (Fig. 1, 3, 6).

**Figure 6.**
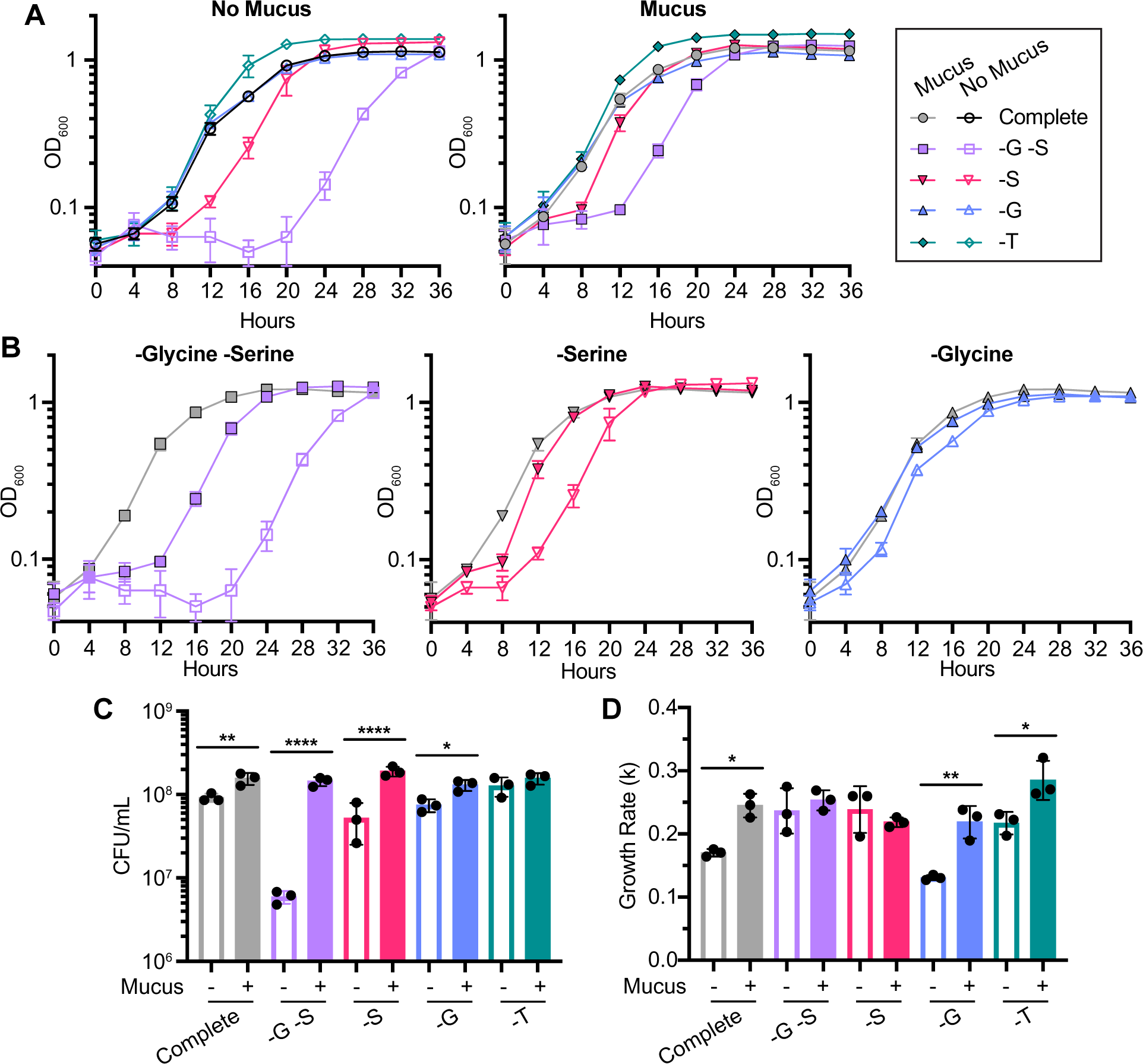
Mucus restores growth in media lacking specific amino acids. **(A)** *C. difficile* growth curves in a modified version of CDMM with defined casamino acids (DCAMM) without mucus (left) and with mucus (right), comparing growth between complete medium and media lacking specific amino acids. **(B)** Growth curves from (A) comparing growth with and without mucus (filled vs. open symbols) for specific conditions: lacking glycine and serine (left), lacking serine only (middle), and lacking glycine only (right). The growth curve from complete medium with mucus is shown on each graph for comparison (grey). (C) Viable cell counts (CFU/mL) at exponential phase. For each pair of media conditions with and without mucus, samples were collected when cultures for either condition first reached exponential phase (OD_600_ ∼0.5), which occurred at: 12 hours for complete, -G, and -T media; 16 hours for -S medium; and 20 hours for -G -S medium. (D) Growth rates during exponential phase. * p < 0.05, ** p < 0.01, *** p < 0.001, **** p < 0.0001 determined by (C, D) one-way ANOVA with Sidak’s test.

Removing threonine from the medium did not affect growth, but addition of mucus to -T media promoted growth overall relative to conditions without mucus, as indicated by slightly faster exits from lag phase (Fig. 6A) and increased growth rates (0.255±0.074 OD_600_/h with mucus vs. 0.217±0.018 without, Fig. 6D). However, we did not observe changes in CFU between conditions with and without mucus in -T media at mid-exponential phase (Fig. 6C).

Conditions lacking both glycine and serine resulted in significant growth defects, particularly a prolonged lag phase (Fig. 6A, 6B). Addition of mucus to -G -S medium enhanced growth (Fig. 6B), yielding 24.3-fold more CFU/mL than -G -S medium without mucus (Fig 6C). Growth rates between the two conditions were similar (Fig. 6D). Overall, a lack of glycine and serine lengthened lag phase, and addition of mucus partially ameliorated this effect (Fig. 6B).

Cultures in -S medium without mucus also took longer to exit lag phase relative to -S medium with mucus (Fig. 6A, 6B). Similar to -G -S conditions, addition of mucus increased growth such that curves appeared more similar to those from complete media with mucus (Fig. 6B). In addition, growth in -S media with mucus yielded 3.65-fold more CFU/mL with mucus versus without (Fig. 6C). In contrast, growth in -G media largely matched that in complete media (Fig. 6A, 6B), with 1.74-fold more CFU/mL and significantly increased growth rates with mucus compared to no mucus (*k* = 0.219±0.026 vs. 0.131±0.004 OD_600_/h) (Fig. 6C, 6D). Together, these results indicate that lack of serine has a greater impact on *C. difficile* growth than glycine in the absence of mucus and suggests mucus serves as a source of serine.

## DISCUSSION

In this study, we demonstrated that mucus derived from primary human IECs enhances *C. difficile* growth and produces a variety of responses in the pathogen, particularly changes in metabolism. These changes emphasize the metabolic plasticity and complexity of *C. difficile* in its adaptive responses to mucus, which may involve differential sensing and transcriptional regulation. Conversely, induction of biofilm formation by *C. difficile* may impact the viscoelastic properties of mucus, which has ramifications for persistence.

Prior work examining the ability of *C. difficile* to utilize mucus as a nutrient source indicated that other bacterial species are needed to liberate moieties from mucus. Our finding that a low concentration of mucus produced a consistent, albeit subtle, increase in growth indicates *C. difficile* can metabolize mucus independently. Indeed, this concentration was even sufficient to elicit dramatic shifts in gene expression. Nonetheless, it remains likely that contributions from other species *in vivo* enhances the use of mucus as a growth substrate.

Distinct properties of the mucus models used may have contributed to these different outcomes. Prior work employed mucus from commercialized sources and immortalized cell lines and applied a purification process resulting in a mono-component mucus containing a single mucin type (MUC2)^20,23^. In contrast, our strategy was to remove free sugars and nutrients while retaining as many features of native mucus as possible. Thus, the mucus we used to supplement media would have retained more of the components and mucin types of native colonic mucus, better reflecting nutrient availability in the colon.

In the presence of mucus, we observed increased expression of genes contributing to CO_2_ fixation via the WLP, and of genes encoding enzymes within the GCS, which is linked to the WLP via methylene-THF (Fig. 7). The WLP may be particularly important later in infection, such as when electron acceptors for Stickland metabolism are low^54^. However, others have reasoned that the WLP remains important under heterotrophic conditions to allow utilization of CO_2_ produced during glycolysis^35^. Metabolic modeling emphasized the importance of the GCS and the conversion of serine to glycine in the presence of mucus (Fig. 7), as predicted fluxes from the GCS and from GHMT indicate that glycine biosynthesis is particularly important. Glycine is a key electron acceptor in Stickland metabolism, where it is reduced to acetylphosphate via GR^33,34^. We observed reduced expression of GR with mucus, suggesting alternative mechanisms for glycine metabolism under the conditions tested.

**Figure 7.**
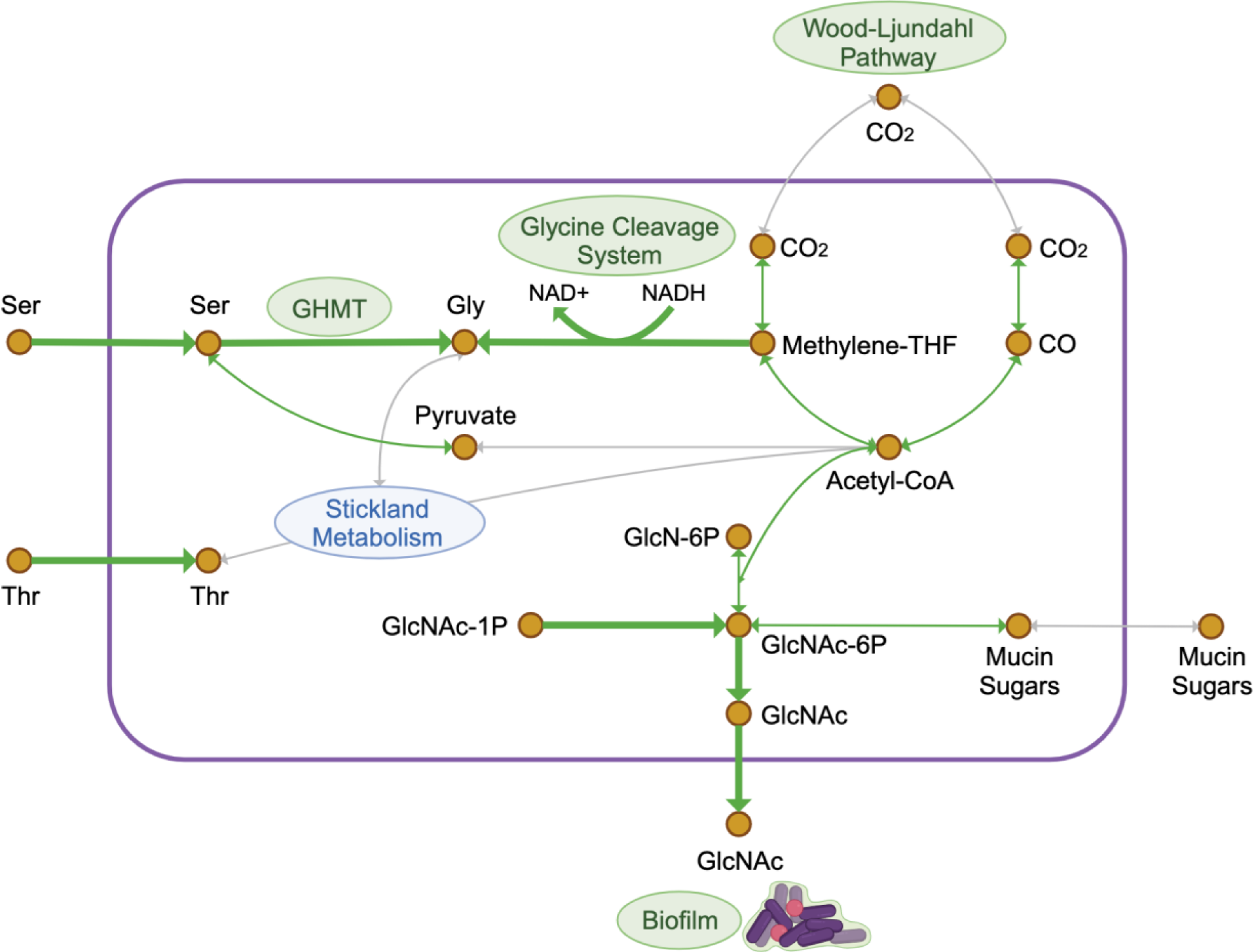
Overview of responses to IEC mucus in *C. difficile*. Simplified diagram of key reactions in the metabolic model and/or identified in the RNA-Seq experiment. Thicker green arrows indicate select reactions determined to be important in Random Forests analysis following contextualization of a *C. difficile* R20291 metabolic model; arrow directions indicate the direction of flux predicted with mucus. Paths within which we identified upregulation of genes corresponding to enzymes are identified with thin green lines. Abbreviations: Ser, serine; Thr, threonine; Gly, glycine; GHMT, glycine hydroxymethyltransferase; Methylene-THF, methylene-tetrahydrofolate; GlcN-6P, glucosamine-6-phosphate; GlcNAc, N-acetylglucosamine; GlcNAc-6P, N-acetylglucosamine-6-phosphate; GlcNAc-1P, N-acetylglucosamine-1-phosphate; CO_2_, carbon dioxide; CO, carbon monoxide. Created with Biorender.com.

Serine also appears to be important during growth in mucus. Metabolic models predicted uptake of serine only when mucus is present and a role for serine in glycine biosynthesis via GHMT. Consistent with this prediction, the growth defect observed in DCAMM lacking serine was improved by adding mucus, suggesting that mucus is a source of serine for *C. difficile*. Importantly, serine deamination is one of the largest contributors to the pyruvate pool in *C. difficile*^36^ (Fig. 7), thus influencing glycolysis, the TCA cycle, and the WLP^34^. In addition, pyruvate fermentation is linked to toxin repression^55,56^, so an increase in serine uptake from mucus has potential ramifications for *C. difficile* pathogenesis. Overall, serine and glycine likely contribute synergistically to the flexibility of *C. difficile* metabolism and its growth.

Experimental data and metabolic modeling suggest that mucus stimulates biofilm formation (Fig. 7). *C. difficile* has been demonstrated to form biofilms on purified MUC2, particularly in co-culture with *Fusobacterium nucleatum*^13^, suggesting *C. difficile* can form biofilms in the human colon. Several biofilm-related genes, such as those involved in GlcNAc biosynthesis^57,58^ and polysaccharide II synthesis and export^59^, were upregulated in the mucus condition (File S2). In line with these data, metabolic modeling predicted the efflux of GlcNAc, a component of biofilm polysaccharides from various organisms^60–63^. Furthermore, when provided alongside pyruvate, GlcNAc promotes biofilm formation^52^. Past work in *B. subtilis* indicated GlcNAc export was enhanced by manipulating features of glucose metabolism^64^; we can thus speculate that alterations to *C. difficile* metabolism may promote the GlcNAc efflux predicted by modeling (Fig. 7). Others have noted increased production of proteins from the WLP and GCS in some biofilm models^65^, indicating that there are several potential connections between *C. difficile* metabolic activity and mucus-induced biofilm formation.

Biophysical and biochemical analyses did not demonstrate substantive breakdown of mucins by *C. difficile* independently, which fits prior evidence that multiple species working in concert are required for efficient degradation of mucins^20,66,67^. Nonetheless, given the repertoire of carbohydrate-active enzymes in *C. difficile* R20291^21^, activity may exist to cleave glycans or other fragments from mucus without producing detectable differences in molecular weight. Indeed, we observed significant upregulation of 31 of the 72 CAZy genes in *C. difficile* (File S2), including enzymes with potential mucolytic activity^20,68,69^.

Many genes related to transcriptional regulation and signal transduction were differentially expressed in response to mucus. In a past study looking at the transcriptional response of *Pseudomonas aeruginosa* to mucus-supplemented media^70^, researchers concluded that selection for specific regulatory and signaling mechanisms likely led to the differential expression of metabolism-related genes they observed with mucus. The differential expression of transcriptional regulators in our dataset could contribute to the phenotypic and metabolic changes we observed with mucus, but further exploration into these mechanisms is needed.

Finally, deleting genes upregulated during growth in mucus, which included genes with predicted roles in sugar uptake or transport, did not lead to growth defects in mucus. Instead, mutants exhibited growth that exceeded that of wildtype. These unexpected growth phenotypes could be related to tradeoffs in adaptation to versus growth in mucus. The identification of many differentially expressed genes related to transcriptional control and signal transduction supports a role for mucus as a stimulus to alter *C. difficile* metabolism, and indeed, mucin glycans have been widely demonstrated to stimulate shifts in microbial gene expression and behavior^71^.

Moreover, it has been posited that a greater capacity to sense multiple nutrients in an environment is metabolically costly, contributing to longer lag phases^72^. Under these assumptions, inefficiencies in sensing, as might be expected for strains lacking transporters^73^, could result in shorter lag phases but incur other long-term fitness costs. Further analysis for gene essentiality may identify mechanisms *C. difficile* relies on in its interactions with mucus.

Overall, our data support a model in which colonic mucus promotes *C. difficile* growth as a source of specific amino acids and a likely source of carbohydrates, which could translate to enhanced colonization *in vivo*. Alterations in expression observed with mucus highlight the metabolic flexibility of *C. difficile*, which is likely a key factor for biofilm formation and persistence in a dynamic host environment^35^. Importantly, biofilms are a main contributor to recurrence^74^, providing an environment protected from antibiotics and from which spores can be generated to promote continued infection^75^. Our work demonstrates that *C. difficile* possesses mechanisms to take advantage of this niche. In better understanding *C. difficile*-mucus interactions, more effective therapeutics to disrupt colonization or promote mucosal immunity can be developed.

## MATERIALS AND METHODS

### Bacterial strains and growth conditions

Table S3 lists strains and plasmids used in this study. *Clostridioides difficile* R20291 was routinely cultured in TY (3% w/v Bacto tryptone, 2% w/v yeast extract, 0.1% w/v thioglycolate)^76^, BHIS (3.7% w/v brain heart infusion, 0.5% w/v yeast extract, 0.1% w/v cysteine)^76^, *C. difficile* minimal medium (CDMM)^29^, or CDMM with defined amino acids (DCAMM, described below and in File S1) at 37°C in an anaerobic chamber (Coy) with an atmosphere of 5% CO_2_, 10% H_2_, and 85% N_2_. *E. coli* strains were grown aerobically at 37°C in lysogeny broth (LB, Miller), and conjugations with *C. difficile* performed anaerobically. Methods for cloning and *C. difficile* mutagenesis, as previously applied^77,78^, are detailed in Text S1. Preparation of *C. difficile* cultures for growth assays are detailed in Text S1.

### Primary human intestinal epithelial cell (IEC) cultures and generation of a mucus layer

All IECs were derived from the transverse colon of a 23-year-old male cadaveric donor. Cultures were maintained in Sato’s Expansion Medium (Sato’s EM)^79^. Transwells (0.4 µm pore, PET, Corning) were coated with 10 µg/mL collagen at least four hours prior to seeding cells in Expansion Medium (EM). EM was replaced every other day until a confluent monolayer formed (5-7 days, or until trans-epithelial electrical resistance measured ≥ 500 ohms x cm^2^). To generate a mucus layer, confluent monolayers were cultured in differentiation medium with 330 ng/mL vasoactive intestinal peptide (DM+VIP) in an air-liquid interface^27,28^. DM+VIP was replaced daily. File S1 contains compositions of cell culture media.

### *Ex vivo* mucus purification

After 4-5 days of differentiation, mucus was removed from the apical surface of the IECs. Epithelia were rinsed with PBS to collect residual mucus. Collected mucus and rinses were stored at −20°C prior to purification. Mucus samples were pooled and filtered using nominal molecular weight limit membranes (Amicon Ultra 3K, Millipore Sigma) to remove free nutrients, metabolites, and contaminants. This purified mucus was suspended in PBS, and total mucus protein concentration was determined using Pierce^TM^ BCA Protein Assay (Thermo Fisher) for standardization^12^. For supplementing media, a relatively low standard concentration of 50 µg/mL mucus was used due to limited availability of IEC-derived mucus.

### *C. difficile*-IEC co-cultures

Mucus-producing IECs (4-5 days in DM+VIP) were used to assess growth of *C. difficile* with or without a mucus barrier. Controls lacking mucus were prepared by removing the mucus layer and rinsing epithelia three times with PBS. PBS was then added to the apical compartment (100 µl, approximately the same volume as mucus). IECs were transferred to an anaerobic chamber and inoculated with *C. difficile* (10^3^ CFU, MOI ∼0.01). To recover *C. difficile* after co-culture, 0.1% DTT in PBS was added to each apical compartment, then the plate was placed on a rocking platform for 20 minutes at room temperature break down mucus. CFU in the apical compartment were enumerated by plating serial dilutions.

### RNA-sequencing

Early stationary phase cultures in TY broth were pelleted and washed with PBS, then inoculated into CDMM with or without 50 µg/mL mucus at an OD_600_ of 0.05. At exponential phase (OD_600_ ∼0.5), cultures were collected and RNA extracted using TriZol as described^80^, followed by purification and DNase treatment (RNeasy kit, RNase-Free DNase Set, Qiagen). RNA was submitted for 150 bp paired-end sequencing on an Illumina HiSeq platform (Azenta Life Sciences). Read processing, including quality assessment, filtering, and mapping steps, were performed using established bioinformatic tools^81–83^. To perform genomic feature counting, we used a prokaryote-specific algorithm, Feature Aggregate Depth Utility (FADU; v1.8)^84^. After obtaining read counts per gene, we used DESeq2 (v1.42.0) in R for differential expression analysis^85^. For Gene Set Enrichment Analysis (GSEA, v4.1.0)^86,87^, we used normalized counts from DESeq2 and gene sets from KEGG. We then used default ranking metrics generated by GSEA to run GSEAPreranked and generate normalized enrichment scores for each gene set. We also narrowed the list of genes applied to GSEAPreranked to only those meeting a differential expression threshold (Benjamini-Hochberg corrected p < 0.05, fold change > 2), using Log_2_ fold change as the ranking metric. Select differentially expressed genes were validated using qRT-PCR as previously described^78,88,89^. Text S1 contains further RNA isolation, read processing, and qRT-PCR details.

### *Ex vivo* mucus preparation for biophysical and biochemical characterization

Mucus was collected and stored at −20°C, then pooled before use to achieve sufficient material. To the pool, CDMM salts (final concentration in mucus 0.5X), trace salts (1X), iron sulfate heptahydrate (1X), and vitamins (1X) (File S1) were added to provide minimally necessary components for *C. difficile* survival. To ensure adequate mucus concentration (>2.5% solids), aliquots were dried and percent solids determined by comparing initial aliquot weight to dry weight. To prepare *C. difficile* inoculums, cultures were grown to exponential phase in CDMM with 50 µg/mL mucus, then pelleted and washed with CDMM salts (1X) and trace salts (1X) (File S1, hereafter 1X CDMM salts). Mucus samples were inoculated with 10^6^ CFU *C. difficile* in 1X CDMM salts at a 1:20 dilution. To prepare controls lacking *C. difficile*, we mock-inoculated mucus with 1X CDMM salts. Samples were collected at inoculation and 24 hours post-inoculation to confirm *C. difficile* viability and for biophysical and biochemical analyses.

### Biophysical and biochemical analysis of *ex vivo* mucus

Biophysical analysis using particle tracking microrheology (PTMR) to determine complex viscosity of *ex vivo* mucus was performed as described^28,90,91^. Biochemical analyses using multiangle laser light scattering (MALLS) to measure mucin molecular weights, radii of gyration, and concentrations in *ex vivo* mucus were also performed as described^28^. Specific details for each technique are in Text S1.

### Biofilm assays

Overnight cultures of *C. difficile* grown in TY broth were pelleted and washed with PBS, then diluted 1:30 in CDMM with or without 50 µg/mL mucus. Cultures were grown to an OD_600_ 0.8-1, normalized toOD_600_ 0.5, and aliquoted into untreated 96-well polystyrene plates. Assay methods were adapted from past work^92–94;^ details are in Text S1.

### Context-specific metabolic modeling

We contextualized a published *C. difficile* R20291 genome-scale metabolic network reconstruction (GENRE) for conditions with and without mucus as described^49–51^, using transcript per million values from the RNA-Seq experiment and the maxfit_contextualize() function in RIPTiDe with default settings. The model was constrained to fit minimal media conditions used (CDMM with or without mucus)^49^. Bray-Curtis dissimilarity, nonmetric multidimensional scaling, and PERMANOVA statistical testing were performed using the vegan R package (v.2.6-4)^50^. To determine reactions important for differentiating conditions, supervised machine learning was performed using the randomForest R package (v. 4.7-1.1)^50^. Differences in predicted fluxes were determined using Wilcoxon rank sum test^50^.

### Growth experiments with defined amino acids minimal medium (DCAMM)

To test the contribution of glycine, serine, or threonine to *C. difficile* growth while keeping conditions consistent with prior experiments, we created a defined casein amino acids minimal medium (DCAMM, File S1). To approximate relative proportions of each amino acid in 10 mg/mL casamino acids (the concentration in CDMM), amino acid content for bovine casein (alpha-S1, alpha-S2, and beta casein subunits, UniProt IDs P02662, P02663, P02666 respectively) was used. All other components of DCAMM were unchanged. *C. difficile* growth curves in each media with or without 50 µg/mL mucus were performed as described in Text S1.

## Data Availability

Unless otherwise noted, all statistical analyses were performed using R (v4.3.2) or GraphPad Prism 10. R, Python, and bash scripts for applying bioinformatic tools are available from GitHub: https://github.com/klfurtado/2024_Cdiff_Mucus_Paper. RNA-Seq reads are available from NCBI Gene Expression Omnibus (<u>GSE254621</u>).

## Supporting information

Supplemental Materia

## ACKNOWLEDGEMENTS

This work was supported by NIH R01-DK120606 to RT and NLA; NIH R01-AI143638 to RT; and NIH P30-DK065988 and Cystic Fibrosis Foundation HILL20Y2-OUT to DBH. NLA has a financial interest in Altis Biosystems. We thank Kimberly Walker for her careful review of the manuscript and Jilarie Santos Santiago for her assistance in maintaining IEC cultures.

