## Supplementary material for "*Clostridioides difficile*-mucus interactions encompass shifts in gene expression, metabolism, and biofilm formation": Tables S1 and S2

**Supplementary Table 1**: Differentially expressed genes related to nutrient acquisition or metabolism. L2FC±SE: $Log_2$ fold change ± standard error.

| ***Fructose and Mannose Metabolism Gene Set (KEGG)*** | | | |
|---|---|---|---|
| **Gene** | **Name, Description, & Details** | **L2FC±SE** | **p-adj** |
| CDR0692 | *gutA*; Glucitol/sorbitol PTS system EIIC component; homologous to CD630 *srlA* | 1.367±0.104 | 0.0000 |
| CDR0693 | *srlE*; Glucitol/sorbitol PTS system EIIB component; homologous to CD630 *srlEa* | 1.498±0.116 | 0.0000 |
| CDR0694 | *srlE'*; Glucitol/sorbitol PTS system EIIB component; homologous to CD630 *srlEb* | 1.493±0.094 | 0.0000 |
| CDR0695 | *srlB*; Glucitol/sorbitol PTS system EIIA component | 1.493±0.114 | 0.0000 |
| CDR0696 | *gutD*; Sorbitol-6-phosphate 2-dehydrogenase; homologous to CD630 *srlD* | 1.176±0.162 | 0.0000 |
| CDR2904 | Mannose PTS system EIIC component | 1.285±0.067 | 0.0000 |
| CDR2175 | L-fuculose-phosphate aldolase | 1.219±0.190 | 0.0000 |
| CDR2898 | *xylA*; Xylose isomerase | 1.040±0.082 | 0.0000 |
| CDR2452 | Fructose PTS system EIIA component | -1.073±0.134 | 0.0000 |
| CDR2976 | Fructose PTS system EIIA component | -1.221±0.229 | 0.0000 |
| ***Wood-Ljungdahl Pathway*** | | | |
| **Gene** | **Name, Description, & Details** | **L2FC±SE** | **p-adj** |
| CDR3174 | *hydN1*; electron transport protein | 0.441±0.070 | 0.0000 |
| CDR3175 | *hydA*; hydrogenase | 0.215±0.066 | 0.0018 |
| CDR3176 | *hydN2*; electron transport protein | 0.361±0.075 | 0.0000 |
| CDR3177 | hypothetical protein; homologous to CD630 *fdh* | 0.489±0.171 | 0.0042 |
| CDR3178 | *fdhD*; formate dehydrogenase accessory protein | 0.214±0.081 | 0.0115 |
| CDR3179 | *fdhF*; formate dehydrogenase | 0.503±0.067 | 0.0000 |
| CDR0648 | conserved hypothetical protein; homologous to CD630 *metV* | 0.270±0.079 | 0.0009 |
| CDR0649 | putative methylenetetrahydrofolate reductase; homologous to CD630 *metF* | 0.270±0.089 | 0.0034 |
| CDR0643 | *cooS*; putative bifunctional carbon monoxide dehydrogenase/acetyl-CoA synthase; homologous to CD630 *acsA* | 0.522±0.074 | 0.0000 |
| CDR0652 | putative carbon monoxide dehydrogenase/acetyl-CoA synthase complex, small subunit, homologous to CD630 *acsE* | 0.516±0.100 | 0.0000 |
| CDR0653 | putative carbon monoxide dehydrogenase/acetyl-CoA synthase complex, alpha subunit, homologous to CD630 *acsC* | 0.853±0.068 | 0.0000 |
| CDR0654 | putative carbon monoxide dehydrogenase/acetyl-CoA synthase complex, methyltransferase subunit, homologous to CD630 *acsE* | 0.575±0.064 | 0.0000 |
| CDR0655 | putative carbon monoxide dehydrogenase/acetyl-CoA synthase complex, beta subunit, homologous to CD630 *acsB* | 0.460±0.069 | 0.0000 |
| CDR0111 | *ptb;* phosphate butyryltransferase | -0.908±0.078 | 0.0000 |
| CDR2571 | putative propanediol utilization protein; homologous to CD630 *pta* | -0.233±0.069 | 0.0010 |
| CDR1012 | *ackA; acetate kinase* | -0.909±0.058 | 0.0000 |
| CDR0112 | *buk*; butyrate kinase | -0.596±0.064 | 0.0000 |
| CDR0915 | *thlA1*; acetyl-CoA acetyltransferase, homologous to CD630 *thlA* | -0.534±0.077 | 0.0000 |
| CDR0914 | *hbd*; 3-hydroxybutyryl-CoA dehydrogenase | -0.449±0.088 | 0.0000 |
| CDR0913 | *crt2*; 3-hydroxybutyryl-CoA dehydratase | -0.516±0.105 | 0.0000 |
| CDR0912 | *etfA2*; electron transfer flavoprotein alpha-subunit | -0.505±0.079 | 0.0000 |
| CDR0911 | *etfB2*; electron transfer flavoprotein beta-subunit | -0.386±0.083 | 0.0000 |
| CDR0910 | *bcd2*; butyryl-CoA dehydrogenase | -0.248±0.067 | 0.0003 |
| CDR2800 | *adhE*; aldehyde-alcohol dehydrogenase | -0.312±0.060 | 0.0000 |
| ***Glycine Cleavage System*** | | | |

| Gene | Name, Description, & Details | L2FC±SE | p-adj |
|---|---|---|---|
| CDR0656 | *gcvH*; putative glycine cleavage system H protein | 0.458±0.086 | 0.0000 |
| CDR0650 | putative carbon monoxide dehydrogenase/acetyl-CoA synthase complex, dihydrolipoyl dehydrogenase subunit; homologous to CD630 *gcvL* | 0.347±0.077 | 0.0000 |
| CDR1556 | *gcvPB*; glycine cleavage system P protein | -0.909±0.077 | 0.0000 |
| CDR1555 | putative bi-functional glycine dehydrogenase/aminomethyl transferase protein, homologous to CD630 *gcvT* | -0.601±0.068 | 0.0000 |
| CDR2615 | *glyA*; putative serine hydroxymethyltransferase | 0.366±0.070 | 0.0000 |
| CDR3082 | *sdaB*; L-serine dehydratase | 0.707±0.067 | 0.0000 |
| ***Proline and Glycine Reductases (Stickland Metabolism)*** | | | |
| **Gene** | **Name, Description, & Details** | **L2FC±SE** | **p-adj** |
| CDR3097 | *prdF*; putative proline racemase | -1.793±0.061 | 0.0000 |
| CDR3098 | conserved hypothetical protein | -2.229±0.064 | 0.0000 |
| CDR3099 | conserved hypothetical protein | -2.180±0.089 | 0.0000 |
| CDR3100 | conserved hypothetical protein | -2.204±0.069 | 0.0000 |
| CDR3101 | *prdB*; proline reductase | -2.245±0.061 | 0.0000 |
| CDR3103 | *prdA*; proline reductase subunit protein | -2.224±0.070 | 0.0000 |
| CDR3104 | *prdR*; sigma-54-dependent transcriptional activator | -0.691±0.083 | 0.0000 |
| CDR3105 | *prdC*; putative electron transfer protein | -1.249±0.083 | 0.0000 |
| CDR2234 | LysR-family regulatory protein | 0.556±0.103 | 0.0000 |
| CDR2235 | putative membrane protein | 0.421±0.143 | 0.0036 |
| CDR2236 | putative Xaa-Pro dipeptidase | 0.258±0.085 | 0.0034 |
| CDR2237 | *grdD*; glycine/sarcosine/betaine reductase complex component C alpha subunit | -3.460±0.070 | 0.0000 |
| CDR2238 | *grdC*; glycine/sarcosine/betaine reductase complex component C beta subunit | -3.306±0.081 | 0.0000 |
| CDR2239 | *grdB*; glycine reductase complex component B gamma subunit | -1.207±0.078 | 0.0000 |
| CDR2240 | *grdA*; glycine/sarcosine/betaine reductase complex component A | -2.292±0.073 | 0.0000 |
| CDR2241 | *grdE*; glycine reductase complex component B alpha and beta subunits | -3.399±0.073 | 0.0000 |
| CDR2242 | *trxA2*; thioredoxin | -3.148±0.101 | 0.0000 |
| CDR2243 | *trxB3*; thioredoxin reductase | -3.155±0.071 | 0.0000 |
| CDR2244 | *grdX*; putative glycine reductase complex component | -2.502±0.085 | 0.0000 |

**Supplementary Table 2**: Differentially expressed genes related to transcriptional regulation or sensing. L2FC±SE: $Log_2$ fold change ± standard error.

| ***Transcriptional Regulators*** | | | |
|---|---|---|---|
| **Gene** | **Name, Description, & Details** | **L2FC±SE** | **p-adj** |
| CDR0508 | TetR-family transcriptional regulator | 2.010±0.072 | 0.0000 |
| CDR1650 | putative transcriptional regulator | 1.670±0.072 | 0.0000 |
| CDR1351 | putative transcriptional regulator | 1.610±0.167 | 0.0000 |
| CDR1936 | GntR-family transcriptional regulator | 1.574±0.098 | 0.0000 |
| CDR1579 | TetR-family transcriptional regulator | 1.541±0.071 | 0.0000 |
| CDR0865 | GntR-family transcriptional regulator | 1.534±0.104 | 0.0000 |
| CDR1467 | MarR-family transcriptional regulator | 1.410±0.077 | 0.0000 |
| CDR0310 | TetR putative transcriptional regulator | 1.367±0.089 | 0.0000 |
| CDR1646 | TetR-family transcriptional regulator | 1.273±0.076 | 0.0000 |
| CDR2050 | putative transcriptional regulator | 1.216±0.140 | 0.0000 |
| CDR0317 | ArsR-family transcriptional regulator | 1.194±0.087 | 0.0000 |
| CDR2553 | AraC-family transcriptional regulator | 1.185±0.085 | 0.0000 |
| CDR1619 | putative transcriptional regulator | 1.171±0.117 | 0.0000 |
| CDR2450 | MerR-family transcriptional regulator | 1.147±0.076 | 0.0000 |
| CDR1950 | putative transcriptional regulator | 1.121±0.142 | 0.0000 |
| CDR3274 | GntR-family transcriptional regulator | 1.106±0.108 | 0.0000 |
| CDR0991 | PadR-family transcriptional regulator | 1.105±0.216 | 0.0000 |
| CDR3067 | MarR-family transcriptional regulator | 1.083±0.095 | 0.0000 |
| CDR1259 | sigma-54 dependent regulatory protein | 1.044±0.092 | 0.0000 |
| CDR1504 | GntR-family transcriptional regulator | 1.026±0.267 | 0.0001 |
| CDR2599 | LysR-family transcriptional regulator | 1.011±0.090 | 0.0000 |
| CDR0506 | TetR-family transcriptional regulator | -1.021±0.085 | 0.0000 |
| CDR3197 | GntR-family transcriptional regulator | -1.041±0.117 | 0.0000 |
| CDR0745 | GntR-family transcriptional regulator | -1.074±0.088 | 0.0000 |
| CDR0817 | GntR-family transcriptional regulator | -1.091±0.076 | 0.0000 |
| CDR0373 | sigma-54-dependent transcriptional regulator | -1.173±0.083 | 0.0000 |
| CDR1311 | GntR-family transcriptional regulator | -1.197±0.118 | 0.0000 |
| CDR2781 | GntR-family transcriptional regulator | -1.313±0.092 | 0.0000 |
| CDR2975 | AraC-family transcriptional regulator | -1.343±0.179 | 0.0000 |
| CDR2929 | *treR*; GntR-family transcriptional regulator | -1.635±0.142 | 0.0000 |
| CDR2847 | putative transcriptional regulator | -1.740±0.119 | 0.0000 |
| ***Sigma Factors*** | | | |
| **Gene** | **Name, Description, & Details** | **L2FC±SE** | **p-adj** |
| CDR1348 | *rpoD2*; RNA polymerase sigma factor RpoD | 1.259±0.136 | 0.0000 |
| CDR0050 | *sigH*;RNA polymerase sigma-H factor | -1.259±0.066 | 0.0000 |
| ***Transcription Antierminators*** | | | |
| **Gene** | **Name, Description, & Details** | **L2FC±SE** | **p-adj** |
| CDR2970 | *bglG2*; transcription antiterminator | 1.469±0.149 | 0.0000 |
| CDR0690 | putative transcription antiterminator | 1.128±0.143 | 0.0000 |
| CDR2222 | *mtlR;* putative transcription antiterminator | 1.036±0.087 | 0.0000 |
| CDR0054 | *nusG*; transcription antitermination protein | -1.098±0.075 | 0.0000 |
| CDR2977 | transcription antiterminator | -1.241±0.155 | 0.0000 |
| ***Two Component Systems*** | | | |
| **Gene** | **Name, Description, & Details** | **L2FC±SE** | **p-adj** |
| CDR1476 | putative two-component histidine kinase | 1.557±0.098 | 0.0000 |
| CDR1569 | two-component sensor histidine kinase | 1.260±0.141 | 0.0000 |
| CDR1568 | two-component response regulator | 1.198±0.140 | 0.0000 |
| CDR2187 | two-component sensor histidine kinase | -1.393±0.079 | 0.0000 |
| CDR0869 | two-component system response regulator | -1.447±0.132 | 0.0000 |

| | | | |
|---|---|---|---|
| CDR2188 | two-component system response regulator | -1.679±0.108 | 0.0000 |
| CDR2020 | two-component sensor histidine kinase | -2.049±0.072 | 0.0000 |
| CDR2021 | two-component system response regulator | -2.372±0.089 | 0.0000 |
| CDR2610 | *hexK*; two-component sensor histidine kinase | -3.268±0.092 | 0.0000 |
| CDR2611 | *hexR;* two-component response regulator | -3.286±0.094 | 0.0000 |
| CDR2206 | two-component sensor histidine kinase | -4.054±0.119 | 0.0000 |
| CDR2205 | two-component response regulator | -5.025±0.122 | 0.0000 |
