## Supplemental Materia for "*Clostridioides difficile*-mucus interactions encompass shifts in gene expression, metabolism, and biofilm formation"

**Text S1.**
**SUPPLEMENTAL METHODS**

**Strain and plasmid construction.** Antibiotics were used as needed for selection or counterselection at the following concentrations: Chloramphenicol (Cm), 20 µg/ml; Ampicillin (Amp) 100 µg/ml; Thiamphenicol (Tm), 10 µg/ml; Kanamycin (Kan), 100 µg/ml; Anhydrotetracycline (Atc), 100 ng/ml. To make in-frame deletions in *C. difficile*, a previously established toxin-antitoxin allelic exchange plasmid, pMSR0, was used (1). Regions of homology (~1 kb) flanking the sequence to be deleted were amplified by PCR. Following digestion of pMSR0 with BamHI, homology arms were inserted into the plasmid using Gibson Assembly. After transformation into *E. coli* DH5$\alpha$, Cm-resistant clones containing the insert were identified by PCR. Insert sequences were confirmed by Sanger sequencing. Plasmids were introduced into *C. difficile* by conjugation with *E. coli* HB101(pRK24) using heat shock as previously described (2). After 24-48 hours, transconjugants were selected on BHIS-TmKan agar, followed by counterselection on BHIS-Atc agar (3, 4). Deletion mutants were idenitified using PCR with primers flanking the homology region, followed by Sanger sequencing to confirm each deletion. Primers used in this study are in Table S4.

**Growth curves.** *C. difficile* cultures were grown approximately 12 hours in TY broth. Cultures were pelleted and washed at least once and resuspended in PBS to remove trace medium. *C. difficile* was added to minimal or defined media with or without 50 µg/mL mucus at a starting optical density at 600nm ($OD_{600}$) of 0.05. Availability of IEC-derived mucus was limited, hence this relatively low concentration of mucus. $OD_{600}$ measurements were taken up to 48 hours. Growth rates were determined by calculating the change in $OD_{600}$ over time during exponential growth using the exponential growth equation in Prism 10, with the exponential growth phase defined by at least three consecutive $OD_{600}$ measurements indicating linear growth. Serial dilutions of each culture were plated at the start of every curve and at exponential and/or stationary phase time points to enumerate CFU/mL and confirm growth.

**RNA isolation and RNA-Seq read processing.** Following RNA extraction with TriZol (5), aqueous layers containing nucleic acids were combined 1:1 with cold 70% ethanol and transferred to columns in a RNeasy kit (Qiagen) for wash and elution steps. On-column DnaseI treatment (Rnase-Free Dnase Set, Qiagen) was performed prior to elution. RNA was stored at -80°C prior to sequencing. After RNA-sequencing, read quality was assessed using FastQC (v0.11.8). Universal sequencing adapter sequences were removed using Trimmomatic (v0.36) (6). Because mucus was derived from human cell cultures, and because PhiX spike-ins are commonly used for quality control in high throughput sequencing workflows, we used Fastq_screen (v0.14.0) to remove any contaminating reads aligning to human mitochondria (Accession No. NC_012920.1) or PhiX (Accession No. NC_001422.1) (7). Trimmed and cleaned reads were aligned to the *C. difficile* R20291 genome (Accession No. FN545816.1) using non-gapping aligner Bowtie2 (v2.4.1) in global mode (8).

**Validation of RNA-Seq results by qRT-PCR**. *C. difficile* broth cultures in CDMM with or without mucus were prepared as described for growth curves. Samples were collected at exponential phase (OD600 ~0.5), and RNA was isolated as described for the RNA-Seq experiment. cDNA was synthesized using a high capacity cDNA reverse transcription kit with random hexamers (Applied Biosystems). cDNA synthesis reactions were also performed without reverse transcriptase to control for genomic DNA contamination. For all reactions, SensiMix SYBR & Fluorescein qPCR reagents (Meridian Biosciences) were used. Primers were used at a final concentration of 500 µM and 2 ng cDNA template were added per reaction. Primer sequences for qRT-PCR are in Table S4. qRT-PCR data were analyzed using $\Delta\Delta$Ct method with *rpoC* as a housekeeping gene as previously described (4, 9, 10). Secondary normalizations were made to the no mucus condition.

**Biophysical and biochemical analysis of *ex vivo* mucus.** To determine the complex viscosity of *ex vivo* mucus, microscopic viscoelastic moduli were quantified using particle tracking microrheology (PTMR) as previously described (11). Fluorescent carboxylated beads (1 µm, ThermoFisher Scientific) were added at a 1:60 dilution at time of inoculation. At each time point, mucus (5 µL) was added to glass slides with a parafilm window and sealed with a glass coverslip. Brownian diffusion of the fluorescent beads was recorded using video microscopy, with motion tracked and quantified using custom Python and MATLAB scripts (TrackPy: https://zenodo-org.libproxy.lib.unc.edu/records/7670439). Viscoelastic moduli were calculated using particle mean squared displacement and the Stokes-Einstein relationship (12, 13). Multiangle laser light scattering (MALLS) was used to measure molecular weights, radii of gyration, and mucin concentrations in *ex vivo* mucus, as described (11). Mucus (10 µl) was diluted 1:100 in 6M guanidinium HCl (Fisher Scientific) to break noncovalent bonds, leaving large mucin molecules. Samples (300 µl) were injected into a 100 µL loop and eluted in light-scattering buffer containing 200 mM NaCl, 10 mM EDTA, and 0.01% $NaN_3$. Samples were run through a size-exclusion chromatography column (Sepharose CL-2B, Cytiva) in series with MALS (DAWN Heleos II, Wyatt Technologies) and differential refractometry (Optilab, Wyatt Technologies). Mucin concentrations were recorded using the differential refractometer.

**Biofilm Assays.** Overnight cultures of *C. difficile* grown in TY broth were pelleted and washed with PBS, then diluted 1:30 in CDMM with or without 50 µg/mL mucus. Cultures were grown to late exponential phase ($OD_{600}$ 0.8-1) then normalized to an $OD_{600}$ of 0.5 before aliquoting into untreated 96-well polystyrene plates. After 24 hours, supernatants were removed, and biomass was fixed with 100% methanol for 20 minutes (14). After fixation, biofilms were stained with 0.1% w/v crystal violet for 30 minutes (15), after which they were gently rinsed with 1X PBS and resolubilized in 4:1 isopropanol acetone, similar to past work (16). To quantify biofilm, absorbance at 570 nm was measured.

### STRAIN AND PRIMER TABLES

**Table S3**: Bacterial strains and plasmids used in this study.

| Lab Notation | Strain/Plasmid Name | Description | Reference |
|---|---|---|---|
| AC472 | *E. coli* DH5α | F- φ80lacZΔM15 Δ(lacZYA-argF)U169 *recA1 endA1 hsdR17*(rk -, mk+) *phoA supE44 thi-1 gyrA96 relA1* λ-*tonA* | Invitrogen (17) |
| RT273 | *C. difficile* R20291 | Ribotype 027 strain (Genbank Accession No. FN545816.1) | (18) |
| RT3118 | *C. difficile* R20291 Δ0453-0455 | Mutant with in-frame deletion of genes corresponding to locus tags CDR20291_0453-0455 | This work |
| RT3148 | *C. difficile* R20291 Δ0693-0696 | Mutant with in-frame deletion of genes corresponding to locus tags CDR20291_0693-0696 | This work |
| RT3120 | *C. difficile* R20291 Δ2495 | Mutant with in-frame deletion of genes corresponding to locus tags CDR20291_2495 | This work |
| RT270 | *E. coli* HB101(pRK24) | *E. coli* strain with plasmid for conjugations with *C. difficile*. $Amp^R$, $Cm^R$ | (19) |
| RT2460 | pMSR0 | Plasmid backbone containing toxin-antitoxin system for allelic exchange in *C. difficile*. $Cm^R$ | (1) |
| RT3318 | pMSR0::0453-0455 | Allelic exchange plasmid containing homology arms flanking 0453-0455 region. $Cm^R$ | This work |
| RT3320 | pMSR0::0693-0696 | Allelic exchange plasmid containing homology arms flanking 0693-0696 region. $Cm^R$ | This work |
| RT3319 | pMSR0::2495 | Allelic exchange plasmid containing homology arms flanking 2495 region. $Cm^R$ | This work |

**Table S4**: Primers used in this study.

| ***Cloning and Mutagenesis:*** | | |
|---|---|---|
| **Lab Notation** | **Primer Name** | **Sequence (5'-3')*** |
| R2743 | pMSR0 Ins Screen F | gtgttatcaattgcactactcatgg |
| R2744 | pMSR0 Ins Screen R | gttgaaccattagctaaggattcag |
| R3418 | 0453-0455 Screen F | ctttttcctgtgtcaatgc |
| R3419 | 0453-0455 Screen R | cttttttgacagtatggcc |
| R3424 | 0453-0455 Gibson UP F | GATTTCTTTCAGTTTC**GGATCC**ggttttccatgtccagg |
| R3425 | 0453-0455 Gibson DN F | gacttataaagttttatcgtgtttttccatgatttcctcccca |
| R3426 | 0453-0455 Gibson UP R | tggggaggaaatcatggaaaaacacgataaaactttataagtc |
| R3427 | 0453-0455 Gibson DN R | GACGTCGACTCTAGA**ggatcc**caaacctatctgccaactc |
| R3651 | 0453-0455 Confirm Deletion F | cattaaacatttttcaccacc |
| R3652 | 0453-0455 Confirm Deletion R | gttttatccttaattaaggacatg |
| R3448 | 0693-0696 Screen F | gaggtaataatatggacccag |
| R3449 | 0693-0696 Screen R | gtagtgaaaatttgtcaggttc |
| R3454 | 0693-0696 Gibson UP F | GATTTCTTTCAGTTTC**GGATCC**gaagagtggcaataggc |
| R3455 | 0693-0696 Gibson DN F | gtgtatagaggaggataaattatatgactggtggacaagttatgtaag |
| R3456 | 0693-0696 Gibson UP R | cttacataacttgtccaccagtcatataatttatcctcctctatacac |
| R3457 | 0696-0696 Gibson DN R | GACGTCGACTCTAGA**ggatcc**gcatgatacagtagcaacg |
| R3549 | 0693-0696 Confirm Deletion F | gcagttagatatttgttagtggg |
| R3550 | 0693-0696 Confirm Deletion R | cttaattacaaaaatgaggctatctc |
| R3648 | 2495 Confirm pMSR0 Ins | cttcttcatccattgcacc |
| R3366 | 2495 Screen F | caagtaattctatagcattcgc |
| R3367 | 2495 Screen R | cagttcttgtcatatcagcac |
| R3372 | 2495 Gibson UP F | GATTTCTTTCAGTTTC**GGATCC**gctcttattattacgaatggag |
| R3373 | 2495 Gibson DN F | cattttatttagctgttttcttttccatcctaatttcatttccc |
| R3374 | 2495 Gibson UP R | gggaaatgaaattaggatggaaaagaaaacagctaaataaaatg |
| R3375 | 2495 Gibson DN R | GACGTCGACTCTAGA**ggatcc**ctttaagttgttccatagcc |
| R3521 | 2495 Confirm Deletion F | cagcatccttaattctctgtgc |
| R3522 | 2495 Confirm Deletion R | ctacttttgcattaaatacggaag |
| ***qRT-PCR:*** | | |
| R3318 | CDR0455 F | gcaattccaacaacttcaccag |
| R3319 | CDR0455 R | agcctctatagtactgttccctatg |
| R3612 | CDR0693 F | aaaggttctggaggatggg |
| R3613 | CDR0693 R | ttactgcagtttctggctttg |
| R3614 | CDR2495 F | cagcgcaaattaaagctcctg |
| R3615 | CDR2495 R | agctaagtctacagcatcaagc |
| R850 | rpoC F | ctagctgctcctatgtctcacatc |
| R851 | rpoC R | ccagtctctcctggatcaacta |

*Restriction enzyme sites are bolded. Capital letters designate Gibson primer tails.

**SUPPLEMENTAL FIGURES**

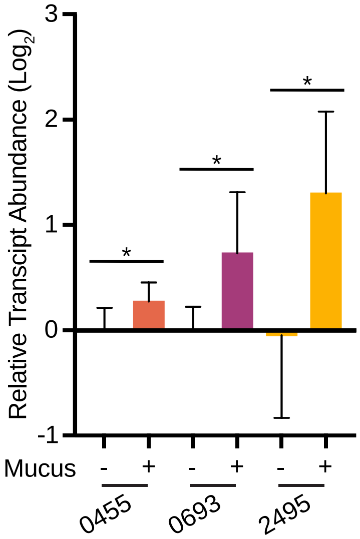


**Figure S1. Confirmation of differential expression of genes of interest.** Relative transcript abundance determined by qRT-PCR for CDR0455, CDR0693, and CDR0696 in broth cultures recapitulating conditions used for RNA-Seq. Mean expression and standard deviations from n=7 or 8 biological replicates shown, combined from 3 independent experiments. *$p < 0.05$, unpaired, two-tailed t-test.

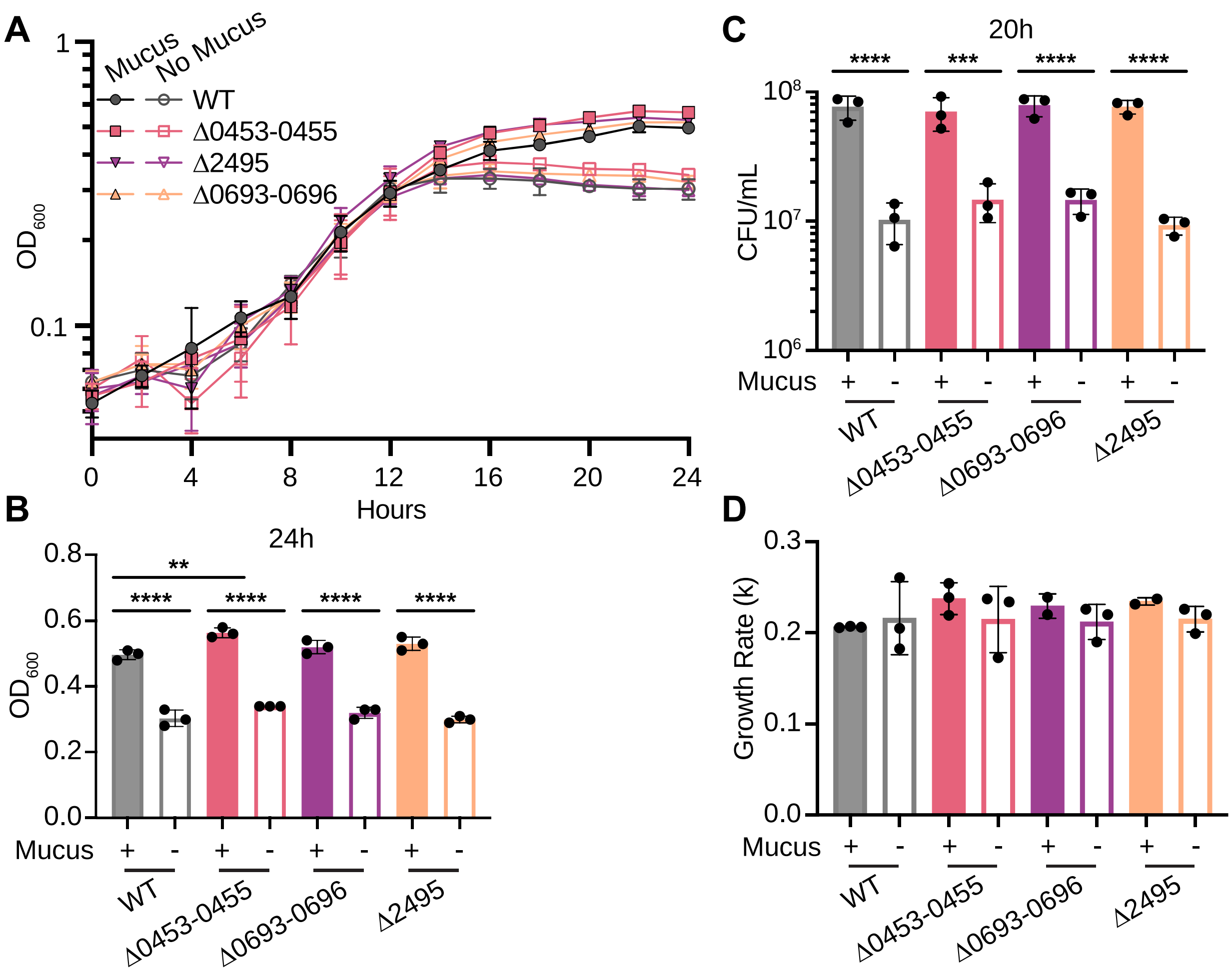


**Figure S2. Effects of deletion of genes and operons upregulated with mucus on growth without glucose. (A)** Growth curves for mutants and wildtype in CDMM lacking glucose with mucus (filled symbols) and without mucus (open symbols). Data are from one representative experiment, n=3. **(B)** Growth rates during exponential phase. **(C)** Viable cell counts expressed as CFU/mL. **(D)** Comparison of $OD_{600}$ values at the final time point. **p < 0.01, ***p < 0.001, ****p < 0.0001, one way ANOVA with Sidak's test.

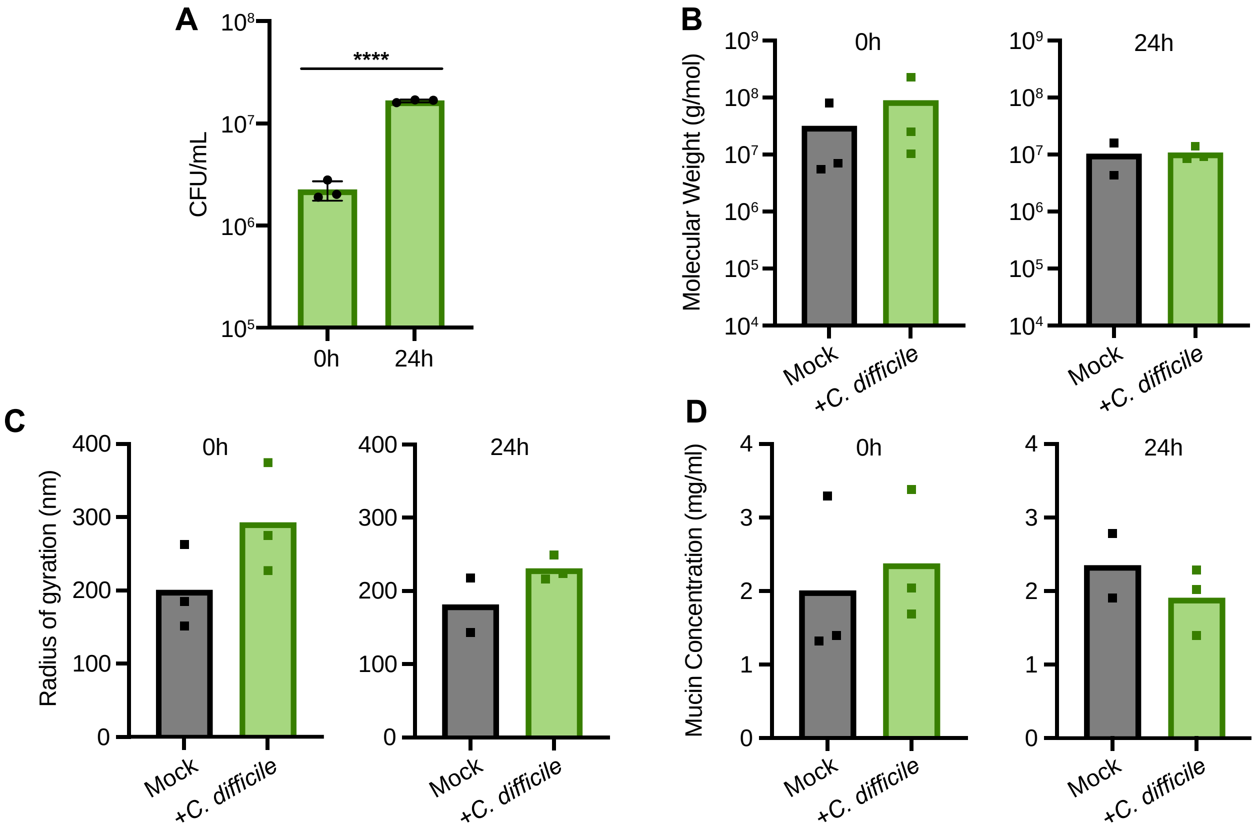


**Figure S3. *C. difficile* viability in *ex vivo* mucus and its effect on mucus biochemical properties. (A)** Viable cell counts for *C. difficile* in *ex vivo* mucus at time of inoculation and after 24 hours, expressed as CFU/mL. ****$p < 0.0001$, unpaired, two-tailed t-test. For biochemical analyses, mucins were separated using size exclusion chromatography and detected by multiangle laser light scattering (MALLS), which measured molecular weights **(B)**, radii of gyration **(C)**, or concentration of mucins **(D)**. For MALLS, each sample was run in technical duplicate or triplicate; means for each sample are shown.
